# No evidence that predictions and attention modulate the first feedforward sweep of cortical information processing

**DOI:** 10.1101/351965

**Authors:** Josipa Alilović, Bart Timmermans, Leon C. Reteig, Simon van Gaal, Heleen A. Slagter

## Abstract

Predictive coding models propose that predictions (stimulus likelihood) reduce sensory signals as early as primary visual cortex (V1), and that attention (stimulus relevance) can modulate these effects. Indeed, both prediction and attention have been shown to modulate V1 activity, albeit with fMRI, which has low temporal resolution. This leaves it unclear whether these effects reflect a modulation of the first feedforward sweep of visual information processing and/or later, feedback-related activity. In two experiments, we used EEG and orthogonally manipulated spatial predictions and attention to address this issue. Although clear top-down biases were found, as reflected in pre-stimulus alpha-band activity, we found no evidence for top-down effects on the earliest visual cortical processing stage (<80ms post-stimulus), as indexed by the amplitude of the C1 ERP component and multivariate pattern analyses. These findings indicate that initial visual afferent activity may be impenetrable to top-down influences by spatial prediction and attention.

## Introduction

Influential predictive coding theories postulate that predictions derived from past experience reduce the magnitude of sensory responses, and that attention can modulate these effects by boosting prediction precision (Friston, 2009; Rao, 2005). Indeed, recent fMRI studies show that predictions based on visual regularities in the environment can modulate neural responses already at the lowest level of the cortical hierarchy, in primary visual cortex (V1) (e.g., Alink, Schwiedrzik, Kohler, Singer, & Muckli, 2010). Moreover, these effects have been shown to depend on attention (e.g., Kok, Rahnev, Jehee, Lau, & De Lange, 2012). For example, Kok et al. (2012) orthogonally manipulated spatial predictions (stimulus likelihood) and attention (stimulus relevance) and found that V1 activity to predicted stimuli was reduced when stimuli were unattended, reflective of reduced prediction error, but enhanced when stimuli were attended, suggestive of heightened weighting of visual evidence by attention. Yet, other studies reported opposing effects of prediction and attention, with prediction and attention, respectively, reducing and enhancing V1 responses (Boynton, 2009; Kok, Jehee, & de Lange, 2012). Since fMRI has low temporal resolution, it is currently still unclear if these effects of prediction observed in V1 reflect modulations of initial feedforward processing, later recurrent (i.e. feedback) processing, or a summation of both. Based on theories of predictive processing one would expect predictions to modulate visual processing as early as V1 (Clark, 2013). Yet, no study so far has shown that predictions can actually modulate initial visual afferent activity.

In the domain of visuospatial attention, the majority of human studies have found no evidence for the notion that attention can modulate the first feedforward sweep of activation, as reflected in the amplitude of the earliest event-related potential (ERP), the C1 (Baumgartner, Graulty, Hillyard, & Pitts, 2018; Bayer et al., 2017; Di Russo et al., 2012; Di Russo, Martinez, & Hillyard, 2003; Martinez et al., 2001; Martínez et al., 1999; Noesselt et al., 2002). This component peaks before 100ms and flips in polarity dependent on whether the stimulus is presented in the upper or lower visual field, suggesting strong contributions of V1 generators (Di Russo et al., 2003; Kelly, Vanegas, Schroeder, & Lalor, 2013). Spatial attention is, on the other hand, robustly associated with modulations of the subsequent visual-evoked P1 component, which reflects extra-striate processing. Accordingly, attention effects observed in V1 in fMRI studies are typically interpreted as driven by feedback from higher visual areas (Di Russo et al., 2003; Martinez et al., 1999, 2001; Noesselt et al., 2002). However, several human M/EEG studies (Kelly et al., 2008; Poghosyan & Ioannides, 2008; Rauss, Pourtois, & Vuilleumier, 2012; Rauss et al., 2009; Slotnick et al., 2002) challenge this conclusion by showing attentional modulations of the C1. For instance, Kelly et al. (2008) showed that, when individual differences in neuroanatomy are taken into account, spatial attention can increase the amplitude of the initial phase of the C1 (50-80ms post-stimulus), which conceivably more selectively indexes V1 activation. Yet, in a direct replication Baumgartner et al. (2018) recently failed to replicate this finding, adding to the controversy of this issue.

Moreover, previous M/EEG work examining how early spatial attention can influence cortical visual processing did not address the possibility that top-down effects might have been absent in the vast majority of previous studies due to the fact that in these studies, stimuli appeared at attended and unattended locations with equal probability (i.e., 50% cue validity; e.g., Baumgartner et al., 2018; Di Russo et al., 2003; Simon P. Kelly et al., 2008). It is conceivable that the exact timing of attention modulation depends on the probability of a stimulus at a given location. Indeed, fMRI work suggests that stimulus-evoked BOLD responses in V1 are largest when a given stimulus is both relevant and more likely (Kok et al., 2012). Thus, the fact that the vast majority of previous studies examining effects of top-down attention on initial visual cortical afferent activity used attention-directing cues with no predictive value may have prevented them from observing effects at the level of C1. Consequently, at present, it remains unclear whether predictions and attention can modulate the first feedforward sweep of cortical visual information processing, and if so, how.

The aim of the current study was to determine if spatial predictions and/or attention can modulate the earliest stage of cortical visual information processing exploiting the high temporal resolution of EEG. In two experiments we orthogonally manipulated stimulus location predictability and relevance (cf. Kok et al., 2012), using the same cueing task and individual C1 titration procedure as Kelly et al. (2008). This allowed us to determine if prediction and attention can modulate the first phase of the C1 (<80ms) and if they do so in interaction. Both attention (Jehee, Brady, & Tong, 2011) and prediction (Kok, Jehee, et al., 2012) have also been associated with sharpening of neural representations in V1 using BOLD fMRI. Therefore, next to examining modulations of activation strength, using multivariate pattern analysis (MVPA) we also investigated how early prediction and attention may modulate visual representations. Lastly, we also explored their effects on pre-stimulus alpha-band activity, indicative of a top-down bias, and on several later ERP components that capture subsequent processing stages.

## Materials and Methods

### Participants

Thirty-two students participated in Experiment 1 (9 males, average age=23.1 years, SD=3.6; age information was not recorded for two participants, but they were between 18 and 40 years old). Fifteen students (6 males, mean age=21 years, SD=2.07) participated in Experiment 2. All participants, recruited from the University of Amsterdam, were right-handed, reported normal or corrected-to-normal vision, and no history of a psychiatric or neurological disorders. Experiment 1 consisted of three EEG sessions, of which the first was used to ensure a robust C1 ERP component in a given individual. Based on this screening, the final set of participants, which participated in all three sessions, consisted of twenty-one participants (6 males, average age= 22.4 years, SD =3.9; age information is not available for two participants). In Experiment 2, two participants were excluded from the final analysis. One participant was excluded due to extremely unmotivated behavior during the second experimental session, and the other due to the absence of a C1 component at both stimulus locations. The study was approved by the ethical committee of the Department of Psychology of the University of Amsterdam. All participants gave their informed consent and received research credit or money (10 euros per hour) for compensation.

### Experimental design and stimuli

All stimuli were generated using Matlab and Psychtoolbox-3 software (Kleiner et al., 2007), and were presented on a 1920 × 1080 pixels BenQ XL2420Z LED monitor at a 120 Hz refresh rate. Stimuli were viewed with a distance of 90 cm from the monitor in all sessions.

### Experiment 1

#### Procedure

The experimental design and tasks were similar to that of Kelly et al. (2008), who previously reported effects of attention on initial visual afferent activity. Specifically, Experiment 1 consisted of three EEG sessions: a ‘probe’ session and two experimental sessions. The probe session served to identify two locations, diagonally opposite to each other, one in the upper and one in the lower visual field, at which for a given participant a reliable C1 could be detected (cf. Kelly et al., 2008). This was done to account for the large variability in V1 anatomy between participants, which may complicate uniform measurement of the C1 at a single electrode or a cluster of electrode sites when the same location is used for stimulus presentation for all participants (Foxe & Simpson, 2002; Kelly et al., 2008; Proverbio, Alice, Del Zotto, & Zani, 2007). These two locations were used in a spatial cuing task in the subsequent experimental sessions. By using diagonally opposite locations, the distance between the attended (cued) and unattended (uncued) location was always equal from fixation and the horizontal and vertical meridians. In the probe session, participants performed a simple target detection task (cf. Kelly et al., 2008), while their brain activity was recorded with EEG. Participants were instructed to fixate on a white cross at the center of the screen, while Gabor patches were flashed briefly in a random order at one of eight locations positioned equidistantly around fixation (see Fig. 1). They had to respond with a left mouse button press only when detecting a target, which was a Gabor patch with a black ring superimposed that appeared on 25% of trials. Participants did not have to respond to non-targets. Because stimuli were presented at each location equally often, it was assumed that attention was distributed evenly across the eight locations.

**Figure 1.**
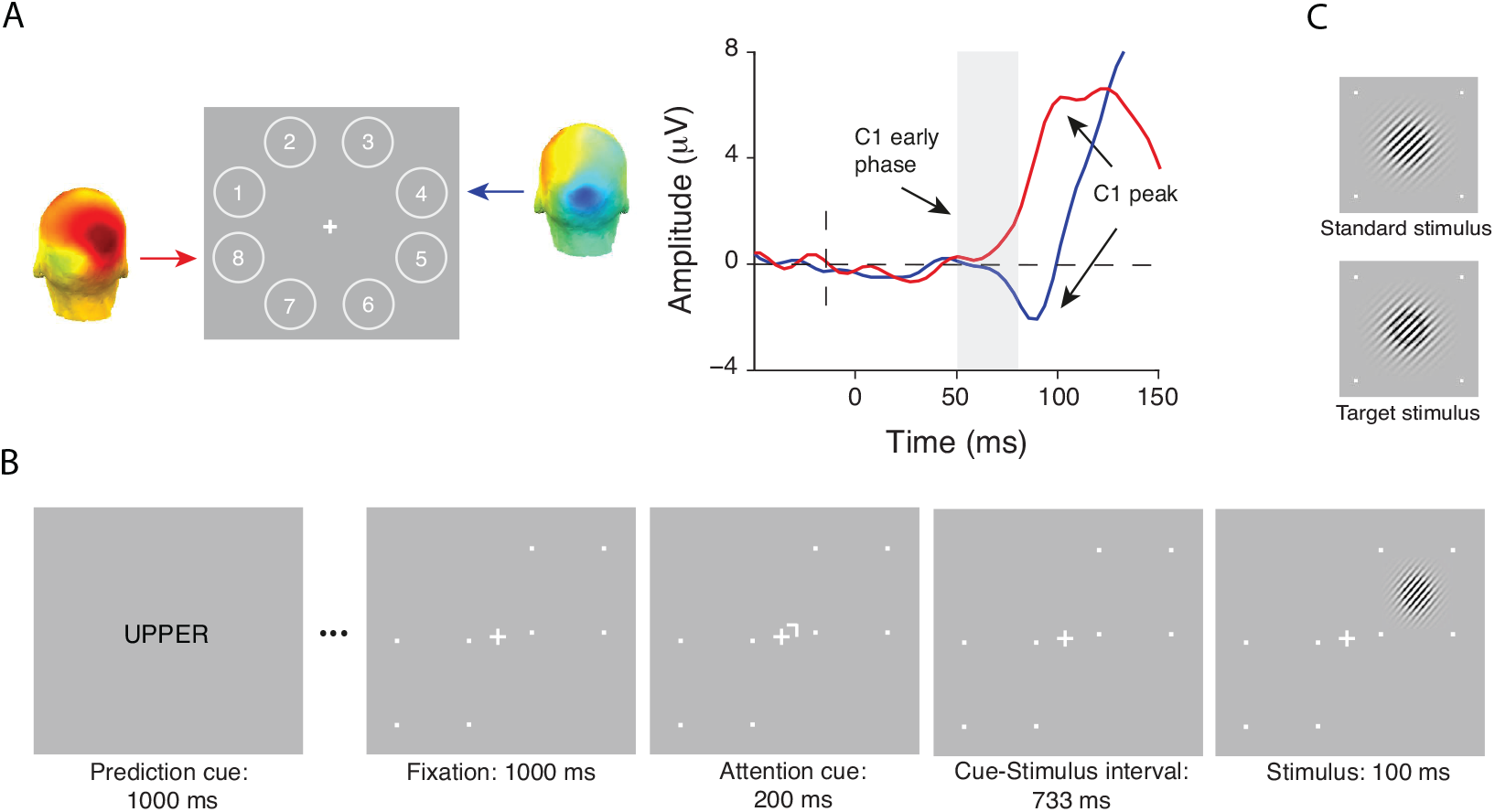
Experimental tasks and stimuli of Experiment 1. (**A**) In the probe session, stimuli were presented at 8 locations around fixation to determine two diagonally opposite locations at which stimuli elicited a robust C1 for a given individual (numbers only shown for display purposes). For a representative subject, shown are the corresponding C1 topographies for an upper location (location 4) and a lower location (location 8) averaged over the 50-80ms post-stimulus period. On the right, the corresponding ERP waveforms are shown. As depicted in the figure, stimuli presented at the upper location elicited a C1 of negative polarity (blue line), whereas stimuli presented at lower location elicited a C1 of positive polarity (red line). (**B**) The spatial cuing task used in the experimental sessions. Each block of the task started with a prediction cue (the word “UPPER”, “LOWER’” or “NEUTRAL”), which signaled the likely location of a stimulus in the upcoming block of 20 trials with 75% (upper or lower cue) or 50% (neutral cue) validity. In each trial, a spatial cue instructed participants to covertly direct their attention to the cued location, which was followed after a fixed delay, by a stimulus, a Gabor patch, at either the cued (attended) or the non-cued (unattended) location. Participants were asked to press a left mouse button if they detected a target, which could only appear at the cued location. Target stimuli appeared on 25% trials and were Gabor patches with a black ring superimposed. The trial sequence shown above is an example of a trial in which a non-target stimulus appears at the location that is both more likely (predicted) and relevant (attended). (**C**) Standard and target stimuli used in Experiment 1.

In two subsequent experimental sessions, administered on separate days, participants performed a visual spatial cueing task virtually identical to the task used by Kelly et al. (2008), (Fig. 1). The crucial difference with Kelly et al. is that we orthogonally manipulated stimulus relevance and probability in the same way as in a previous fMRI study that observed modulations of BOLD activity in V1 by spatial attention and prediction (Kok et al., 2012). This allowed us to study the interactive effects of attention and prediction on initial visual afferent activity. On each trial, a centrally presented attention cue instructed participants which location (the upper or lower) to covertly attend to. This cue was followed by a stimulus, a Gabor patch, at either the cued or the uncued location. Participants had to press a mouse button when detecting a Gabor patch with a black ring superimposed on it (i.e., the target) at the cued (relevant) location. Targets could never appear at the uncued location, i.e., the uncued location was never relevant. The probability of a stimulus appearing at a given location was manipulated in a block-by-block fashion (cf. Kok et al., 2012). At the beginning of each block of trials, a prediction cue informed participants about the likelihood that a stimulus would appear at the upper or lower location in that block. In different blocks, for a given location, this likelihood could be high (75%), neutral (50%), or low (25%). This way, crucially, in a given trial, a stimulus could be attended and predicted, attended and non-predicted (in the “neutral” blocks), attended and unpredicted, unattended and predicted, unattended and non-predicted, or unattended and unpredicted. Participants were instructed to maintain fixation on the center of the screen at all times.

#### Design and stimuli

Non-target Gabor patches in all sessions had a spatial frequency of 6 cycles per degree, a diameter of 1° at half-contrast, and 45° orientation. Targets in all sessions were Gabor patches with a black ring superimposed. The black ring had a radius of 0.4° from the center to outer edge of the ring and thickness of 0.11°.

In the probe session, stimuli appeared randomly at 1 of 8 locations in an annulus of 4° from fixation, and a target stimulus was presented on 11% of the trials (cf. Kelly et al., 2008; Fig. 1). Stimuli were each presented in the middle of an octant (starting with location 1 in the upper visual field at a polar angle of 157.5°, location 2 at 112.5°, and so on). The probe session started with a practice block consisting of 160 trials, followed by 10 blocks of 360 trials each interleaved with self-timed breaks. After every third block, typically a longer break was taken to ensure enough rest throughout the session. To encourage sustained engagement of participants during the probe session, we adaptively controlled task difficulty using a staircase procedure by changing the luminance of the black ring that defined targets (cf. Kelly et al., 2008). Ten difficulty levels varied in a range from a 2% to a 47% reduction in brightness (with steps of 4.5%). Each participant started with difficulty level 4. After two hits with no misses in between, the difficulty level increased by one level. The difficulty level decreased after two false alarms with no misses in between, or after a single miss. After each block, the difficulty level was again reset to level 4.

In two subsequent experimental sessions, participants performed the spatial cuing task while we recorded their brain activity with EEG and monitored fixation using eye tracking (Fig. 1). Each block of 20 trials started with a centrally presented prediction cue that indicated likely location of a stimulus in that block of trials. Prediction cues were words: ‘UPPER’ (75% probability of target appearing at the upper location (i.e., 25% chance of appearing at the lower location)), ‘LOWER’ (75% probability of target appearing at the lower location) or ‘NEUTRAL’ (50% probability of target appearing at either the upper or lower location) presented for 1000ms (cf. Kok et al., 2012). After the prediction cue, a white fixation cross (0.3° in length and 0.12° in width) was shown at the center of the screen for another 1000ms. Each trial in a block started with a centrally presented attention cue, a small white rotated L-shape (line elements were 0.15° in length and 0.06° in width) pointing towards a specific location that participants needed to attend covertly. The attention cue was presented for 200ms at 0.4° eccentricity from the central fixation cross, and pointed to the upper or lower location with equal probability. The direction of the attention cue was randomized within a block of trials so that both locations were equally often relevant in a block. Locations were marked with 4 white small squares outlining a 2.75° × 2.75° area on the screen. A stimulus appeared on the screen 733ms after attention cue offset at 4° eccentricity (cf. Kelly et al., 2008). The inter-trial interval was jittered between 1000 and 1500ms. The difficulty of the spatial cueing task was adjusted online and titrated to 75% correct by adaptively changing the luminance of a black ring on target Gabor stimuli, which appeared on 25% of trials. In comparison to difficulty levels employed in the probe session, here we used more fine-grained step sizes to adjust the task difficulty. There were 40 difficulty levels, again in a range from 2% to 47% reduction in brightness. The starting difficulty level for the first experimental session was the mean difficulty level reached in the probe session. Difficulty level was calculated and adjusted if necessary at 3 points in the experiment (in every forced break; see below). The difficulty would increase 4 levels when mean accuracy was above 90%, and for 2 levels if accuracy was higher than 80%. When accuracy was between 75 and 80%, the difficulty level increased one level, and when it fell between 70 and 75%, it decreased one level. Task difficulty decreased for 2 levels and 4 levels if the performance was below 70% and 60%, respectively. Difficulty levels were adjusted automatically in every forced break, but the experimenter could overwrite this if necessary, for instance, if the false alarm rate was high and hit rate therefore inflated.

Each experimental session consisted of 2008 trials divided into 16 runs. Each run contained 6 blocks of 21 trials, 2 of each prediction condition (“UPPER”, “LOWER”, “NEUTRAL”). Randomization and counterbalancing were done for 502 trials at a time to prevent trials of the same condition to be overrepresented in a certain period of the task. Eight “NEUTRAL” blocks had one trial less due to rounding. After every fourth run there was a longer forced break, and participants could take a shorter, self-timed break in between the other blocks or runs. In every break, participants received written feedback about their performance (percentage of hits, average reaction time and false alarm rate if it was higher than 15%), and a verbal warning if they had broken fixation excessively. They also received written feedback on the computer screen if they had too many false alarms, and were encouraged to keep their performance up to high levels. Before the start of the experiment, every participant practiced two upper, two lower and two neutral prediction blocks.

### Experiment 2

#### Procedure

In Experiment 2, we aimed to determine if we could replicate the findings from Experiment 1 using a further optimized design. Specifically, in Experiment 2, in order to increase the signal-to-noise ratio of neural signal used to test for the effects of attention and prediction, we used large-scale, high-contrast V1-tuned stimuli (Fig. 8) that have been shown to elicit large C1’s (e.g., see Pourtois, Rauss, Vuilleumier, & Schwartz, 2008). Second, subjects performed an orientation discrimination task on the stimuli, which likely relies on V1 processing. Third, stimuli were only presented in the upper field to avoid overlap between the C1 and subsequent P1 component. While the C1 generated by lower field stimuli peaks over lateral posterior electrodes, like the P1, the C1 generated by upper field stimuli peaks over midline posterior electrodes, allowing for a better separation of the two components (Qu & Ding, 2018). The task and procedure were otherwise similar to Experiment 1, except for some additional changes that we detail in the below.

We expected that the V1-tuned stimuli used in Experiment 2 would elicit a detectable C1 component at the scalp in the majority of subjects. Therefore, we skipped the probe session in Experiment 2. Participants thus came to the lab twice for an experimental session in which they performed a spatial cueing task for approximately 120 minutes, while their brain activity was recorded with EEG and eye fixation was monitored with eye tracking. In the first experimental session, they first performed a detection task to titrate the initial difficulty level of the subsequent spatial cueing task. Each task was preceded by detailed instructions explaining the task and a short practice block to familiarize participants with the task.

As in Experiment 1, in the spatial cueing task, spatial attention and prediction were manipulated independently. The to-be-attended (i.e., task relevant) location was cued on a trial-by-trial basis, while location likelihood (25%, 50%, 75%) was varied block-wise. Thus as in Experiment 1, a stimulus could be attended and predicted, attended and non-predicted, attended and unpredicted, unattended and predicted, unattended and non-predicted, or unattended and unpredicted. As in Experiment 1, a response was required only when a target stimulus was presented at the cued location, while standard stimuli did not require a response. A target stimulus could never appear at the uncued location.

#### Design and stimuli

In each trial, following a fixation period of 1000ms, an attention-directing cue indicated which of two locations in the upper field was relevant. The upper-left location was at a polar angle of 135° at 5° eccentricity, and the upper-right at 45° at 5° eccentricity. In Experiment 2, the cue used to direct attention was non-spatial, and consisted of the central fixation cross turning blue or red indicating the left or right location, respectively, as task-relevant for that trial. We used a symbolic cue in Experiment 2, because the cue used in Experiment 1 had a spatial component, which may have exogenously instantiated an attentional bias in the direction of the cued location (the tip of the L shape was pointing to the to-be-attended location). The attention cue was presented for 200ms, and followed by a 833 millisecond-long cue-stimulus interval, after which a stimulus was presented at either the cued or the uncued location (see Fig. 8). In order to boost the strength of stimulus-driven activation in V1, stimuli used in this experiment were high-contrast textures consisting of 7 × 7 white line elements on a black background, all oriented horizontally, except for 3 vertically-oriented line elements in the center of the texture stimulus (4^th^ row and 3^rd^, 4^th^ and 5^th^ column-elements). With respect to the background, these vertical line elements formed a foreground region in the center of the stimulus. The position of each stimulus line was jittered across trials by adding a vertical and horizontal offset that varied between 0 and 0.017° in order to minimize adaptation effects. The entire texture stimulus was 4.75° × 4.75° in size. Each line element was 0.42° × 0.03° and spaced 0.07° apart (see Fig. 8).

Standard and target stimuli differed only in the orientation of the foreground region. The foreground line elements on a standard stimulus formed a 90° orientation contrast with respect to horizontally oriented background lines. Foreground regions on target stimuli formed an orientation contrast different (higher or lower) than 90° (within the bounds of 0 to 180°). The magnitude of the difference between the orientation of the foreground region on a target stimulus and the foreground region on the standard stimulus defined task difficulty. Task difficulty was adaptively changed after every break (self-timed or forced), targeting t=0.5, where t was the difference between the hit and false alarm rate. If it was higher than 0.5, the difficulty of the task increased, i.e., the difference of the foreground orientation between standard and target stimuli decreased by 1°. The difficulty level remained the same for t=0.5, and decreased with one level if the performance dropped below t=0.5, corresponding to an increase in the difference of the foreground orientation contrast of 1°. At each difficulty level, the foreground region on a target stimulus was tilted towards the left (>90°) and towards the right (<90°) equally often.

The starting difficulty level of the spatial cuing task in the first session was determined first for each participant in a separate detection task. This simple detection task consisted of 2 blocks of 200 trials each, divided by a self-timed break. The task and procedure were similar to the spatial cueing task, except that location relevance and location predictability were not manipulated (cf. the probe session in Experiment 1). Stimuli appeared in the upper-left or upper-right quadrant with equal probability. Participants were instructed to press the mouse button only when they detected a target stimulus (texture stimuli with the central foreground region forming the orientation contrast different than 90° with respect to background), while maintaining fixation. Target stimuli appeared on 20% of trials at either location. Four possible target difficulty levels occurred equally likely throughout the task. The difference between target foreground regions with respect to standard foreground regions changed with a step size of 2°. For each of the four difficulty levels (i.e., the highest difficulty level was +/-2° + 90°), the difference between the hit and false alarm rate was computed. The difficulty level at which the performance was closest to t=0.5 was selected as the starting difficulty level of the subsequent spatial cuing task. The difficulty level that a subject reached in the last block of the first experimental session was taken as the starting difficulty level for the second experimental session.

### Data acquisition and preprocessing

#### Eye tracking

Eye movements were recorded using a Tobii X120 infrared eye tracker (120 Hz sample rate) and monitored online throughout each session in both Experiments. A standard 9-point calibration was performed at the start and after every 4 blocks. If the eye position fell outside of a circle with radius of 1.5° around fixation for more than 100ms, the central fixation cross would turn from white to grey, indicating to participants that their eyes were not on fixation and that they had to fixate their gaze at the central fixation cross again.

#### EEG recordings and preprocessing

EEG data, digitized at 512 Hz, were continuously recorded in all sessions in both Experiments using an ActiveTwo system (BioSemi, Amsterdam, the Netherlands), from 64 scalp electrodes placed according to the 10/20 system, four electrooculographic (EOG) electrodes placed above and below, and to the side of the eyes, and two external electrodes attached to each earlobe. EEG data were offline referenced to the average activity recorded at the earlobes, resampled to 265Hz, and filtered using a 50 Hz notch filter and then high-pass filtered at 0.1 Hz, and low-pass filtered at 45 Hz. The continuous data were subsequently epoched from −2.1 to 2.1 seconds around stimulus presentation and baseline corrected to the average activity between −80ms to 0ms pre-stimulus (cf. Kelly et al., 2008). Epochs with EMG artifacts or eye blinks that coincided with stimulus or attention cue presentation were rejected based on visual inspection. Extremely noisy or broken channels were reinterpolated. Remaining eye blink artifacts were removed by decomposing the EEG data into independent sources of brain activity using an Independent Component Analysis (ICA), and removing eye blink components from the data for each subject individually. Preprocessing was done using the EEGLAB toolbox (Delorme & Makeig, 2004) for Matlab (The MathWorks, Inc. Natick, MA, USA) and custom-written Matlab scripts.

### Analyses

Analyses, unless reported otherwise, were done using the EEGLAB toolbox (Delorme & Makeig, 2004) for Matlab (The MathWorks, Inc. Natick, MA, USA), custom-written Matlab scripts (time-frequency analyses) and SPSS (repeated-measures ANOVA and paired sample t-test).

### Experiment 1

#### Eye-tracking

Eye tracking data were analyzed offline to determine for each trial, if eye deviation from fixation was greater than 1.5 degrees for at least 50ms in a −500 to 500ms interval around stimulus presentation, or if eye tracking data was missing for more than 100ms in the same time window. These trials were excluded from behavioral and EEG data analyses. To ensure that the mean gaze deviation across participants did not differ between conditions (e.g., varied as a function of location relevance or location likelihood) at the time of stimulus presentation, we entered eye position values in the x and y direction averaged across −200 to 100ms locked to the stimulus presentation into separate repeated measures ANOVAs with the within-subject factors Attended Location (upper, lower) and Predicted Location (predicted, non-predicted, unpredicted). To enable direct comparison of deviations in the horizontal direction, values in the x-direction were multiplied by −1 for the subset of subjects who had their upper and lower locations in the right and left visual field respectively. Thus, their eye tracking data were transformed, as if, in each subject, the upper location was in the left visual field and the bottom location in the right visual field.

#### The spatial cueing task: behavior

Behavioral analyses were conducted to ensure that our stimulus predictability manipulation in the spatial cueing task impacted behavioral performance. Specifically, we statistically evaluated the difference in average reaction times, accuracy (percentage of correct target detections) and d’ (target sensitivity) between three prediction conditions, collapsed across the upper and lower field conditions, with a repeated measures ANOVA with Prediction (P, NP, UP) as a within-subject factor. d’, a sensitivity index based on signal detection theory (SDT) (Stanislaw & Todorov, 1999), was computed as Z(hit rate)-Z(false alarm rate).

#### The probe session EEG data

Following the procedure described in Kelly et al. (2008), the probe task EEG data was analyzed in order to identify two spatial locations, one in the upper and one in the lower field, diagonal to each other (e.g., upper left location 1 and lower right location 5 in Figure 1 a), where stimuli elicited a robust C1 ERP component. To this end, for each subject separately, we computed ERP waveforms to non-target stimuli for each of the 8 locations separately and inspected the waveforms for the presence of a C1. Following the same procedure as Kelly et al. (2008) and Baumgartner et al. (2018), we defined the C1 based on a combination of component timing, scalp topography, and polarity information. Specifically, the C1 is characterized by 1) an onset around 50ms, 2) a rise above baseline before 80ms and peak before 100ms over posterior scalp regions, and 3) a positive polarity for lower-field stimuli and a negative polarity for upper-field stimuli (Di Russo, Martínez, Sereno, Pitzalis, & Hillyard, 2002; Kelly et al., 2008). Based on these characteristics, a pair of diagonally opposite locations showing a clear C1 was selected for each subject. These were used as stimulus locations in the spatial cuing task in the subsequent two experimental sessions. Out of thirty-two participants tested in the probe session, eleven participants were excluded from further testing, as they did not exhibit a clearly identifiable C1 at two diagonally opposite probe locations.

#### ERP analyses: C1 component

We created ERPs time-locked to non-target stimuli, separately for upper and lower-field stimuli, per condition. One participant’s data did not yield an identifiable C1 component for lower-field stimuli, due to which we excluded his/her data from the C1 analyses. Based on the offline analysis of the eye tracking data, only trials in which the eyes were within 1.5° from fixation were included in the ERPs.

To address our main question, if prediction and attention independently or in interaction can modulate the first feedforward sweep of visual cortical activity, we conducted a 3-way repeated measures ANOVA with Attention (A, UA), Prediction (P, NP, UP) and Field (upper, lower) as within-subject factors, and average voltage values in 50-80ms time window at C1 peak channels as the dependent variable.

To determine possible effects of attention and prediction on the later phase of the C1, which more likely also reflects contributions from extra-striate sources, we repeated the same analyses, but now with C1 peak amplitude as the dependent variable. C1 peak amplitude was determined as follows. Based on the condition-average upper and lower-field ERPs, for each subject separately, we determined at which electrode and latency the amplitude of the C1 was most positive (for lower field stimuli) or negative (for upper field stimuli) within a time window of 50-100ms post-stimulus. The obtained C1 parameters (two peak amplitudes and peak latencies per subject, one for each field) were then used to quantify C1 peak amplitude separately for each condition of interest. On average, the C1 peaked at 93 ms for upper, and at 100ms for lower-field stimuli. Average C1 amplitude over +/-1 sample around the peak sample (~12ms) was then used as the dependent variable in a three-way repeated measures ANOVA to test for the presence of top-down modulations of the later phase of the C1. The repeated measures ANOVA examined the independent and interactive effects of attention and prediction and included the within-subject factors Attention (A, UA), Prediction (P, NP, UP), and Field (upper, lower). In all repeated measures ANOVAs, the polarity of C1 amplitude to upper-field stimuli was inverted so that all values were positive and therefore directly comparable to amplitudes of lower-field stimuli. For all repeated measures ANOVA analyses, here and in following sections, whenever Mauchly’s test suggested a violation of sphericity, we report Geenhouse-Geisser corrected p-values.

In case of non-significant effects of prediction and/or attention on the initial or later phase of the C1, we performed Bayesian statistics using JASP (JASP Team, 2018) software to determine the strength of evidence for the null hypothesis of no effect (Masson, 2011; Wagenmakers et al., 2018). Generally speaking, Bayesian statistics allows the quantification of the probability of hypotheses for and against the absence of effects, i.e. H0 and H1, given the observed data p(H |D). These probabilities can be statistically compared and expressed as a Bayes factor (BF01), which indicates the posterior probability of the H0 over H1 (Masson, 2011). The higher the value of the Bayes factor BF_01_, the stronger the evidence in favor of H_0_ being true. Here, we used terminology for interpreting Bayes factors suggested by Jeffreys (1961), and labeled a BF_01_ from 1 to 3 as anecdotal evidence in favor of H_0_, values from 3 to 10 as substantial, and those above 10 as strong evidence in favor of H0. To evaluate the main and interaction effects of interest, we conducted a Bayesian repeated measures ANOVA with the same within-subject factors to determine the strength of evidence in favor of the H_0_. In case we needed to quantify evidence for an interaction effect, we computed inclusion Bayes factor (BFinclusion) across matched models, which is the ratio between the sum of posterior model probabilities P(M |data) of all models that contain the interaction effect of interest, but no interactions with the interaction effect of interest, and the sum of posterior model probabilities of all the models included in the numerator term but without the interaction of interest. This factor thus indicates the extent to which data supports the inclusion of the interaction effect, taking all relevant models into account. For the sake of consistency in interpreting the BFinclusion in line with the BF01, we inverted (1/ BFinclusion) this factor such that it indicates the evidence in favor of H0 (Wagenmakers et al., 2018) in accordance with terminology suggested by Jeffreys (1961).

#### EEG multivariate analysis: Top-down effects on spatial representations

Both attention (Jehee, Brady, & Tong, 2011) and prediction (Kok, Jehee, et al., 2012) have also been associated with sharpening of neural representations in V1 using BOLD fMRI. Therefore, next to examining modulations of activation strength using univariate ERP analyses (described above), using multivariate pattern analysis (MVPA), we also investigated how early attention and prediction may modulate sensory representations, as reflected in the pattern of EEG activity across electrodes. This multivariate approach may also be more sensitive in picking up weak top-down modulations of activation strength than the analytic approach by Kelly et al. (2008) of only looking at the C1 peak electrode that we followed in our ERP analysis, as this approach takes into account neural activity at many scalp electrodes, as well as inter-individual differences in those pattern (Slagter, Alilović, & van Gaal, 2018). In order to examine the effects of attention and prediction on the representational content of neural activity, as reflected in distribution of neural activity across the scalp we used the ADAM toolbox (Fahrenfort, Leeuwen, Olivers, & Hogendoorn, 2017) to train a linear discriminant classifier to distinguish the patterns of activity between conditions using the raw EEG signal measured at all electrodes (features for classification). Specifically, we tested if patterns of neural activity differ between attended vs. unattended, and predicted vs. unpredicted conditions. We used a 10-fold cross-validation to evaluate classification performance. Raw EEG data was divided into 10 folds (information about the order of trials was removed). Next, a classifier was trained on 90% of the data to classify between stimulus classes, and then tested on the remaining 10% of the data. The training and testing procedure was repeated ten times, so that each time a different portion of data was used for training and testing to avoid circularity (see Kriegeskorte, Simmons, Bellgowan, & Baker, 2010). For each subject, classification accuracy was calculated as the average number of correct condition classifications, first averaged across conditions, and then across 10 folds. This was done for each sample of the EEG signal, which resulted in vector of classification accuracies over time. Classification accuracies were tested using a one-sample two-sided t-test to evaluate if accuracies differed significantly from chance. Intervals of significant decoding were corrected for multiple comparisons using group-wise cluster-based permutation testing, as described in Maris & Oostenveld (2007) and implemented in the ADAM toolbox (Fahrenfort et al., 2017). In this procedure, a sum of t-values in a cluster of temporally adjacent significant time points (p<0.05) in the observed data is computed. This sum is compared to the sum of t-values in a cluster of significant data points obtained under random permutation. Random permutation and computation of a sum of t-values under random permutation is repeated 1000 times. The p-value used to evaluate the significance of a cluster in the observed data is the number of times the sum of t-values under random permutation exceeded that of the observed sum, divided by the number of iterations.

#### Time-frequency analysis: Top-down effects on pre-stimulus alpha activity

Attention and prediction have also been associated with changes in pre-stimulus alpha oscillatory activity (Horschig, Jensen, van Schouwenburg, Cools, & Bonnefond, 2014; Kelly, Lalor, Reill, & Foxe, 2006; Sauseng et al., 2005; Worden, Foxe, Wang, & Simpson, 2000), suggesting that these top-down factors can bias visual processing in advance. To assure that participants indeed used attention and prediction cues to adaptively track stimulus likelihood and shift their attention to cued locations in advance, here we performed a complex Morlet wavelet decomposition of raw EEG data to obtain the time-frequency representation (Mike X Cohen, 2014). The wavelet convolution was performed in the frequency domain, such that the power spectrum of EEG signal (obtained using the fast Fourier transform) was multiplied by the power spectrum of Morlet wavelets. Morlet wavelets were computed by point-wise multiplication of a complex sine wave with a Gaussian window: *e^i2πtf^ e^-t^2^/2σ^2^^* (where *t* is time, *f* is frequency, which increased from 2 to 40 Hz in 30 logarithmically spaced steps, and *σ* the width of the Gaussian, which increased logarithmically from 3 to 8 in the same number of steps). This was done on a single-trial level. Based on the resulting complex signal, the power at each frequency band and time point was computed as the squared magnitude of the result of the convolution: *real*[*z*(*t*)]^2^ + *imaginary* [*z*(*t*)]^2^ (Cohen & Donner, 2013; Cohen, 2014). Power values were baselined to average pre-stimulus power between −1133 to −983ms (i.e., 200 to 50ms before attention cue onset) at each frequency band using a decibel (dB) transformation: *dB power* = 10 *x log_10_*(*power_t_/power_baseline_*) (Cohen & Donner, 2013). To examine the effects of spatial prediction and attention, per condition separately, trial-averaged alpha power values (8-12 Hz) were computed for two pairs of parieto-occipital electrodes, PO4/PO8 and PO3/PO7 in a −500 to −100ms pre-stimulus time-window, where anticipatory effects were expected to be most pronounced (Kelly et al., 2006; Sauseng et al., 2005; Worden et al., 2000). To test if prediction and attention induced alpha power asymmetry between electrode sites contralateral and ipsilateral to the predicted and to-be-attended locations, respectively, these values were submitted to a repeated measures ANOVA with the within-subject factors Attended Location (upper, lower), Predicted Location (P, NP, UP), and Hemisphere (contralateral, ipsilateral). Note that which electrode pair for a given subject and to-be-attended location was considered contralateral or ipsilateral depended on whether the to-be-attended location in the e.g., upper visual field was in the left or the right hemifield. Significant effects were followed up by repeated measures ANOVAs and paired sample t-tests.

#### ERP analyses: Top-down effects on later ERP components

A large body of work has shown that visual processing after 100ms is susceptible to top-down modulation (e.g., Di Russo et al., 2003; Lasaponara et al., 2017; Marzecová, Widmann, Sanmiguel, Kotz, & Schröger, 2017; Noesselt et al., 2002). Therefore, to confirm longer-latency activity modulations, in a set of secondary analyses, we also examined how attention and prediction, separately and/or in interaction, modulated stimulus processing over time, after 100ms. Specifically, we examined their effects on the amplitude of the visual-evoked P1 and N1 components, as well as of the later P3a and P3b components. Consistent with previous studies (e.g., Di Russo et al., 2003; Noesselt et al., 2002), the group- and condition-average P1 and N1 components peaked over lateral occipitoparietal scalp sites. Two pairs of lateral occipitoparietal electrodes (PO4/PO8 and PO3/PO7) were hence used to calculate the amplitude of the P1 and N1 components contralateral and ipsilateral to the stimulus location, separately per condition of interest. The largest positive voltage value in 100-150ms interval, and the largest voltage negativity within 150-200ms were selected to determine the latency of the P1 and N1 peaks, respectively, for each subject separately. Time windows for peak picking were based on visual inspection of the group- and condition-average ERPs. The latencies of the contralateral and ipsilateral P1 and N1 peaks were largely consistent with previous studies. The contralateral P1 peaked at 137ms for upper and at 117ms for lower-field stimuli, and at 145ms for upper and 141ms for lower-field stimuli over ipsilateral sites. The N1 peaked contralaterally at 180ms and 172ms for upper and lower-field stimuli, respectively. The ipsilateral N1 peak was measured at 200ms for upper, and at 181 for lower-field stimuli. Average P1 and N1 amplitude values +/-1 sample around the peak sample were entered into separate repeated measures ANOVAs with four within-subject factors: Attention (A, UA), Prediction (P, UP, NP), Hemisphere (contralateral, ipsilateral), and Field (upper, lower). Significant effects that included the factor(s) Attention and/or Prediction were followed up by paired t-tests.

We also examined effects of prediction and attention on the later P3a and P3b components, which are consistently shown to be modulated by stimulus relevance and probability (e.g., Friedman, Cycowicz, & Gaeta, 2001; Marzecová et al., 2017; Polich, 2007). To this end, based on the condition-average data, collapsed across the upper and lower location, we first determined the peak latency of the P3a over fronto-central electrodes (FCz, Fz) in the 300-400ms time window, and the peak latency of the P3b over parieto-central electrodes (POz, Pz) in the 350-450ms time window. The P3a peaked at 360ms for upper field, and at 348ms for lower field stimuli. The P3b peak was identified at 387ms for upper and at 395ms for lower field stimuli. Mean peak amplitude values (peak -/+ 12 samples around the peak, i.e., ~100ms) were then calculated for each condition separately and averaged across electrodes. These values were entered into separate repeated measures ANOVAs with factors Attention (A, UA) and Prediction (P, NP, UP). Significant effects in all repeated measures ANOVAs were followed-up by paired sample t-tests.

### Experiment 2

#### Eye tracking

Due to a malfunctioning eye tracker, only 5 participants had full eye tracking datasets available in Experiment 2, and their data were analyzed identically to the eye tracking data in Experiment 1 (see above). For the remaining 8 participants, eye-tracking data was missing for more than 15% of trials (collapsed across two sessions). For these participants, we manually inspected the HEOG channel for horizontal eye movements and removed trials with eye movement activity. To exclude possible condition differences in eye position on sensory-evoked ERPs, we statistically compared average x-coordinates (horizontal eye movements), y-coordinates (vertical eye movements), and HEOG recorded voltages in −200 to 100ms interval in separate repeated measures ANOVAs with Attended Location (upper, lower) and Predicted Location (P, UP, NP) as within-subject factors. Following the empirical work of Mangun & Hillyard (1991), we also estimated deviations from fixation in degrees of visual angle based on HEOG voltages measured after cues directing attention to the left and right location, in the −200 to 100ms interval around stimulus presentation.

#### Data acquisition, preprocessing and statistical analysis

Data acquisition procedure and preprocessing steps were comparable to those in Experiment 1. ERP and MVPA analyses were identical to those in Experiment 1. As we were specifically interested in top-down effects on the first feedforward sweep of activation, replication ERP analyses only focused on the C1. The C1 component was again assessed from the signal recorded from an individually determined C1 peak channel at a peak latency determined based on the condition-average data separately for left and right upper field stimuli. The C1 component peaked at 86 and 84ms for left and right upper field stimuli, respectively.

## Results

### Experiment 1

#### Behavior

A repeated measures ANOVA revealed that stimulus predictability modulated the speed of responses to target stimuli (F_2,18_=4.617, p=.005, see Fig. 2A). As in Kok et al. (2012), whose manipulation of location predictability we adopted, participants were significantly faster in detecting target stimuli that occurred at predicted (P=443.3ms, SD=37.6) compared to non-predicted (NP=449.8ms, SD=40.7, t_19_=-2.48, p=.023) and unpredicted (UP=460.2ms, SD=45.9, t_19_=-3.12, p=.006) locations. They were also significantly faster in detecting targets at non-predicted in comparison to unpredicted locations (t_19_=2.83, p=.011). As in Kok et al. (2012), accuracy was not affected by stimulus predictability (F_2,18_=0.43, p=.487; P=76.5, SD=3.9; NP=77.1, SD=4.9; UP=78, SD=8.3) (Fig. 2B). Target sensitivity, as indexed by d’, was also not affected by stimulus predictability (F_2,18_= 1.81, p=.19; d’ P=2.27, SD=0.5; d’ NP=2.39, SD=0.6; d’ UP=2.28, SD=0.6). These results confirm that predictions influenced stimulus processing, as participants were fastest at detecting targets presented at the most likely location. Participants on average accurately detected 74.5% (SD=1.61) of the targets in the probe task and 77.4% (SD=4.8) of the targets in the spatial cueing task, which is comparable to the performance level reported by Kelly et al. (2008) (80.7%, SD=3.3), and was expected given that we titrated performance to 75%. Since target stimuli only appeared at cued (i.e., attended) locations, we could only determine effects of stimulus location predictability, not location relevance, on performance.

**Figure 2.**
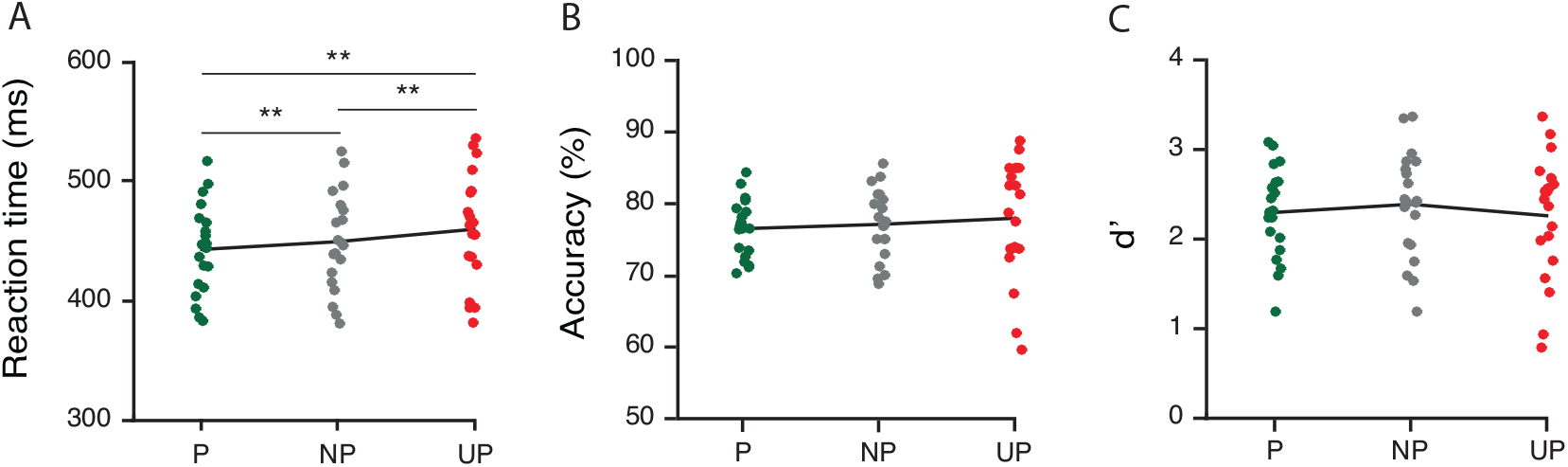
Effects of prediction on behavioral performance in Experiment 1. (**A**) Speed of responses decreased linearly with increasing stimulus predictability: Participants responded significantly faster to predicted than to non-predicted and unpredicted stimuli, as well as to non-predicted compared to unpredicted stimuli (**p<0.01), confirming that our prediction manipulation was successful. Prediction did not significantly influence accuracy (**B**) or d’ (**C**).

#### Early C1 modulations

Our main question was whether prediction and attention, independently or in interaction, may modulate the first feedforward sweep of visual cortical activity, as indicated by the early phase of the C1. Results from a repeated-measures ANOVA addressing this question did not reveal any evidence for top-down modulations: the main effects of Attention (F_1,18_=0.079, p=.781) and Prediction (F_2,17_=0.608, p=.556) were not significant, neither was their interaction (F_2,17_=0.173, p=.685). This was further supported by results of a Bayesian repeated measures ANOVA, which provided substantial to strong evidence for the null hypotheses against a main effect of Attention (B_01_=6.6), a main effect of Prediction (B_01_ = 18.1). Furthermore, inclusion Bayes factor across matched models provided substantial support against an interaction between Attention and Prediction (BF_inclusion_=9.1). The classical repeated measures ANOVA further revealed that the C1 amplitude in the early phase was significantly higher for stimuli presented in the upper vs. lower visual field (main effect of Field: F_1,18_=9.383, p=.007), but the factor of Field did not interact with Attention (F_1,18_=2.489, p=0.132), Prediction (F_1,18_=1.219, p=0.32), nor did it modulate their interaction (F_2,17_=2.479, p=.114). Thus, in contrast to the notion that predictions can modulate afferent activity in V1, we found no evidence for top-down modulation of the early phase (50-80ms) of the C1.

#### C1 peak modulations

Inspection of Figure 3 revealed that the C1 component might be modulated by top-down factors to a greater extent slightly later in time, around its peak. Therefore, we also examined if prediction and/or attention might modulate the peak of the C1. A repeated measures ANOVA also revealed no significant effect of Attention on the later phase of the C1 (F_1,18_=0.742, p=.400). Yet, a significant interaction between Attention and Field (F_2,17_=6.925, p=.017) suggested that attention may have modulated the later phase of the C1 differentially at upper vs. lower locations. This was confirmed post-hoc: the C1 peak was significant larger for stimuli presented at attended vs. unattended locations only when stimuli were presented in lower visual field (t_18_=2.314, p=.033), and not when they were presented in the upper visual field (t_19_=-0.515, p=.612). These findings conceivably reflect differential overlap from the P1 attention effect, that one would expect for lower visual field stimuli only, as in contrast to upper visual field stimuli, which generally elicit a C1 that is maximal over midline electrodes, lower visual field stimuli elicit a C1 that is typically maximal over the same lateral posterior scalp regions as the P1 (Baumgartner et al., 2018; Di Russo et al., 2012, 2003; Kelly et al., 2008; Martinez et al., 2001; Martínez et al., 1999).

**Figure 3.**
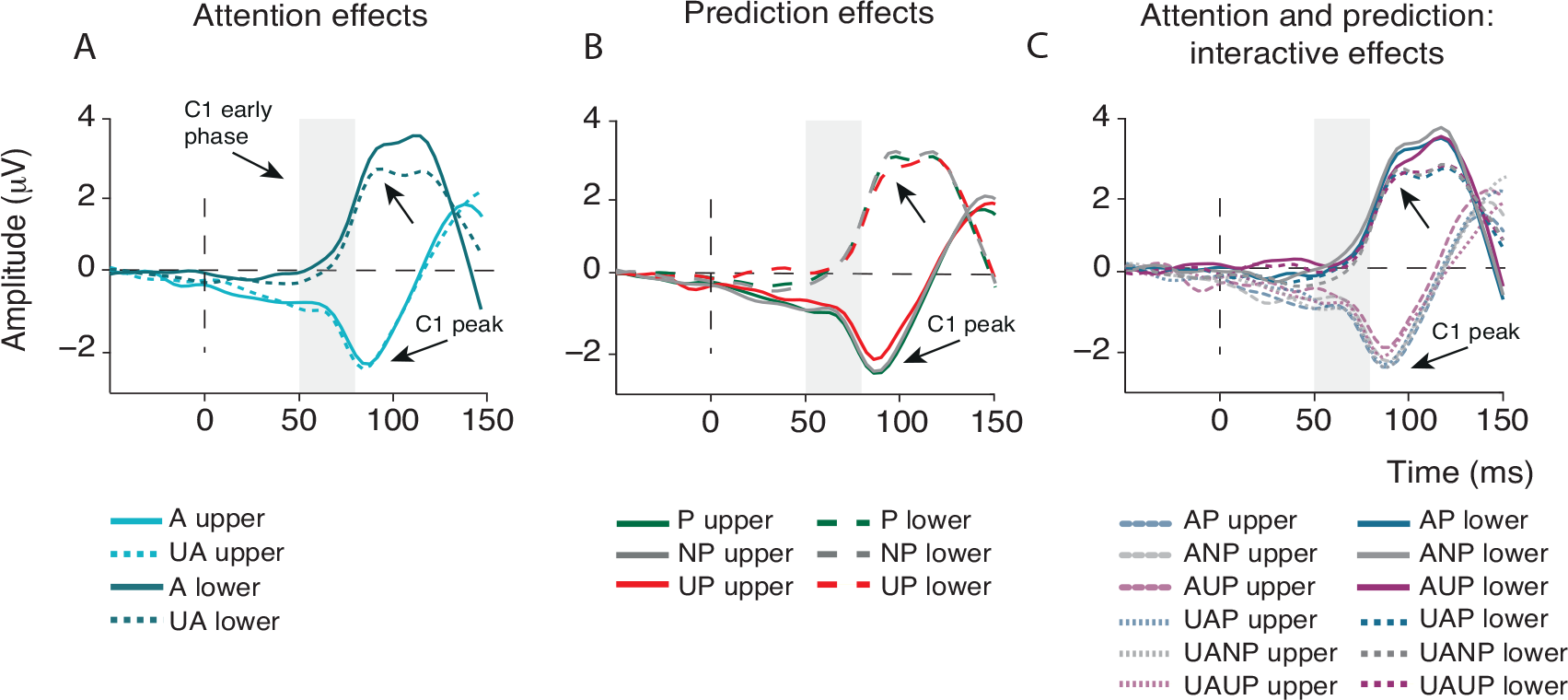
Effects of prediction and attention on the first feedforward sweep of cortical information processing. Shown are grand-average ERPs measured at individually determined C1 peak electrodes, separately for upper and lower visual field stimuli and locked to stimulus onset. (**A**) Attention (collapsed across prediction conditions), did not modulate the early phase of the C1. The C1 peak, slightly later in time, was significantly larger for attended (A) versus unattended (UA) stimuli, but only for stimuli presented in the lower field, likely reflecting overlap from the P1 attention effect at lateral posterior scalp regions. (**B**) Prediction (collapsed across attention conditions) also did not modulate the early phase of the C1 (50-80ms post-stimulus), but did influence the amplitude of the C1 peak. Specifically, the C1 peak in the unpredicted (UP) condition was significantly lower in amplitude than the C1 peak in the non-predicted (NP) and predicted (P) conditions. This is contrary to what one would expect based on predictive processing accounts that postulate that unpredicted stimuli should elicit greater sensory activity (i.e., prediction errors). Moreover, this finding was not supported by Bayesian analyses, which provided stronger evidence for the hypothesis that prediction does not modulate C1 peak amplitude. (C) Attention and prediction in interaction did not modulate the early phase of the C1 or its peak amplitude.

Furthermore, we found that the C1 peak was modulated by stimulus location likelihood, as suggested by a significant main effect of Prediction (F**2,17**=3.743, p=.045). However, contrary to the notion of prediction-based suppression of sensory processing (e.g. Alink et al., 2010; Kok, Jehee, et al., 2012; Kok, Rahnev, et al., 2012) this effect was driven by a significantly larger C1 peak to predicted than to unpredicted stimuli (t_18_=2.4, p=.027), and by a significantly larger C1 peak to non-predicted than to unpredicted stimuli (t_18_=-2.330, p=.032). The difference between predicted and non-predicted C1 peaks did not reach significance (t_18_=.038, p=.970) (see Fig. 3B). The prediction effect was not modulated by the Field (F_2,17_=0.255, p=.778). Finally, attention and prediction in interaction did not modulate the C1 peak amplitude (F_2,17_=0.706, p=.508).

In contrast to the results of the classical repeated measures ANOVA, a Bayesian repeated measures ANOVA yielded strong evidence in favor of the null hypothesis against an effect of Prediction on the later phase of the C1. Specifically, the Bayes factor indicated that the data were 10.9 more likely under the null hypothesis, constituting strong evidence against a main effect of Prediction. Moreover, in line with the classical analysis, the Bayesian analysis suggested substantial evidence for the null hypotheses of no effect of Attention (BF_01_=4.9) and strong evidence for no interaction between Attention and Prediction (BF_inclusion_=10.8). Finally, evidence against a Field by Attention interaction was only anecdotal (BF_inclusion_=2.04), which is in line with the result obtained by using the frequentist approach. Thus, while the classical (frequentist) repeated measures ANOVA suggested that predictions may modulate the later phase of the C1, this was not supported by our Bayesian analysis, which provided strong evidence for the absence of an effect of prediction. Thus, we found no evidence that prediction and/or attention can modulate the early phase of the C1, and mixed evidence for an effect of prediction on the later phase of the C1 in a direction opposite to what one would expect based on predictive processing theories in which predictions are proposed to reduce visual responses (i.e., prediction errors) (e.g. Alink et al., 2010; Friston, 2009).

#### Top-down effects on neural representations

Using multivariate decoding analyses, we next examined potential effects of attention and prediction on representational content, as both attention (Jehee, Brady, & Tong, 2011) and prediction (Kok, Jehee, et al., 2012) have been associated with sharpening of neural representations in V1 using BOLD fMRI. Using a backward decoding model, we obtained a time course of decoding accuracies, indicative of when in time precisely attention and prediction began to modulate neural activity patterns. Fig. 4 shows classification accuracies over time for attended versus unattended (Fig. 4A) and predicted versus unpredicted stimuli (Fig. 4B). The classifier was able to discriminate between attended and unattended conditions with above-chance accuracy from approximately 133 to 992ms post-stimulus (two-tailed cluster p-value <.001 after 1000 iterations). Predictions modulated patterns of neural activity slightly later in time, from approximately 242ms until about 648ms post-stimulus, as revealed by significant decoding (predicted vs. unpredicted) (two-tailed cluster p-value<.001 after 1000 iterations). These multivariate results corroborate and extend the univariate ERP results, and support the conclusion that spatial attention and prediction did not modulate visual information processing before 80ms.

**Figure 4.**
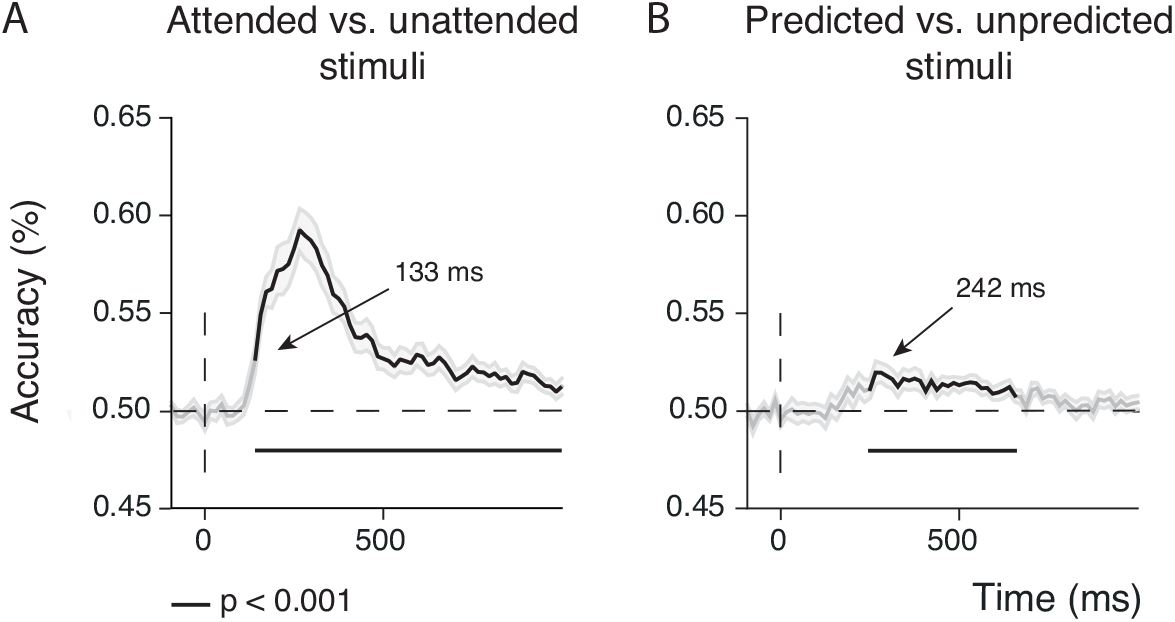
Effects of attention and prediction on the sharpness of stimulus location representations. (**a**) Classification accuracy for attended versus unattended stimuli collapsed across upper and lower locations and prediction conditions. Attention modulated patterns of neural activity between 130 – 990ms post-stimulus (black lines mark two-tailed cluster p-value <.001 after 1000 iterations). (**B**) Classification accuracy for predicted versus unpredicted stimuli collapsed across upper and lower locations and attention conditions. Predictions modulated neural activity patterns between 242 – 650ms (black lines mark two-tailed cluster p-value <.001 after 1000 iterations). Shaded areas on both figures are +/-s.e.m. These multivariate findings corroborate the univariate ERP findings, as they do not reveal any effects of attention and prediction before 80ms.

#### Pre-stimulus alpha power modulations

Given the absence of top-down modulations of visual activity before 80ms, we next examined if attention and prediction modulated prestimulus baseline activity, as indexed by pre-stimulus alpha-band oscillatory activity. By examining attention- and prediction-related changes in pre-stimulus alpha power, we wanted to ensure that subjects in fact directed their attention in advance to the cued location and that location likelihood was used to predict upcoming stimuli in advance. Previous studies have robustly related spatial attention (albeit confounded with prediction as attended stimuli were also more likely than unattended stimuli) with greater alpha activity over ipsilateral versus contralateral posterior scalp regions, in line with the notion that attention can bias sensory regions in advance to favor processing of task-relevant over irrelevant information (Worden et al., 2000). There is also initial evidence to suggest that predictions can modulate pre-stimulus alpha activity (Horschig et al., 2014).

Indeed, although attention did not modulate the stimulus-evoked C1, attention did modulate pre-stimulus alpha-band activity, as suggested by a significant main effect of Hemisphere (F_1,19_=25.470, p<.001). This main effect captured the expected pattern of relatively greater alpha activity over ipsilateral vs. contralateral posterior scalp regions (contralateral power=-0.63, SD=0.66, ipsilateral power=-0.28, SD=0.54) (see Fig. 5A). This asymmetry in pre-stimulus alpha power was not significantly affected by whether attention was directed to the upper or the lower location (Attended Location x Hemisphere interaction: F_1,19_=3.491, p=.077), although total alpha power was significantly higher when upper (upper locations=-0.34, SD=0.54) compared to lower locations (lower locations=-0.57, SD=0.7; main effect Attended Location: F_1,19_=5.518, p=.03) were attended. This analysis confirms that participants covertly directed their attention in advance towards the cued location.

The main effect of Predicted Location (F_2,18_=0.433, p=.591) and the interaction between Predicted Location and Hemisphere (F_2,18_=0.689, p=.445) were not significant. However, a significant Attended Location x Predicted Location x Hemisphere interaction (F_2,18_=9.027, p=.002) suggested that prediction had an effect on pre-stimulus alpha asymmetry that was dependent on the to-be-attended location. Indeed, a post-hoc analysis revealed that, when lower locations were cued, the interaction between Predicted Location and Hemisphere was significant (F_2,18_=5.33, p=.019), while this was not the case when upper locations were cued (F_2,18_=0.568, p=.510). A paired sample t-test further revealed that prestimulus alpha power was significantly greater over ipsilateral vs. contralateral posterior scalp regions when lower locations were cued (i.e., attended) and predicted (t_19_=-3.15, p=.005, predicted contralateral=-0.72, SD=0.85; predicted ipsilateral=-0.38, SD=0.65), but not, for instance, when two locations were equally likely (t_19_=-1.308, p=.207, non-predicted contralateral=-0.67, SD=0.8; non-predicted ipsilateral=-0.52, SD=0.57). These results suggest that when the likelihood that a stimulus would occur at the attended lower visual field location was high, alpha power exhibited the characteristic lateralization in particular for predicted locations (see Fig. 5C). These results suggest that predictions and attention may interact to bias visual regions in advance, however here, only when lower field locations were task-relevant.

**Figure 5.**
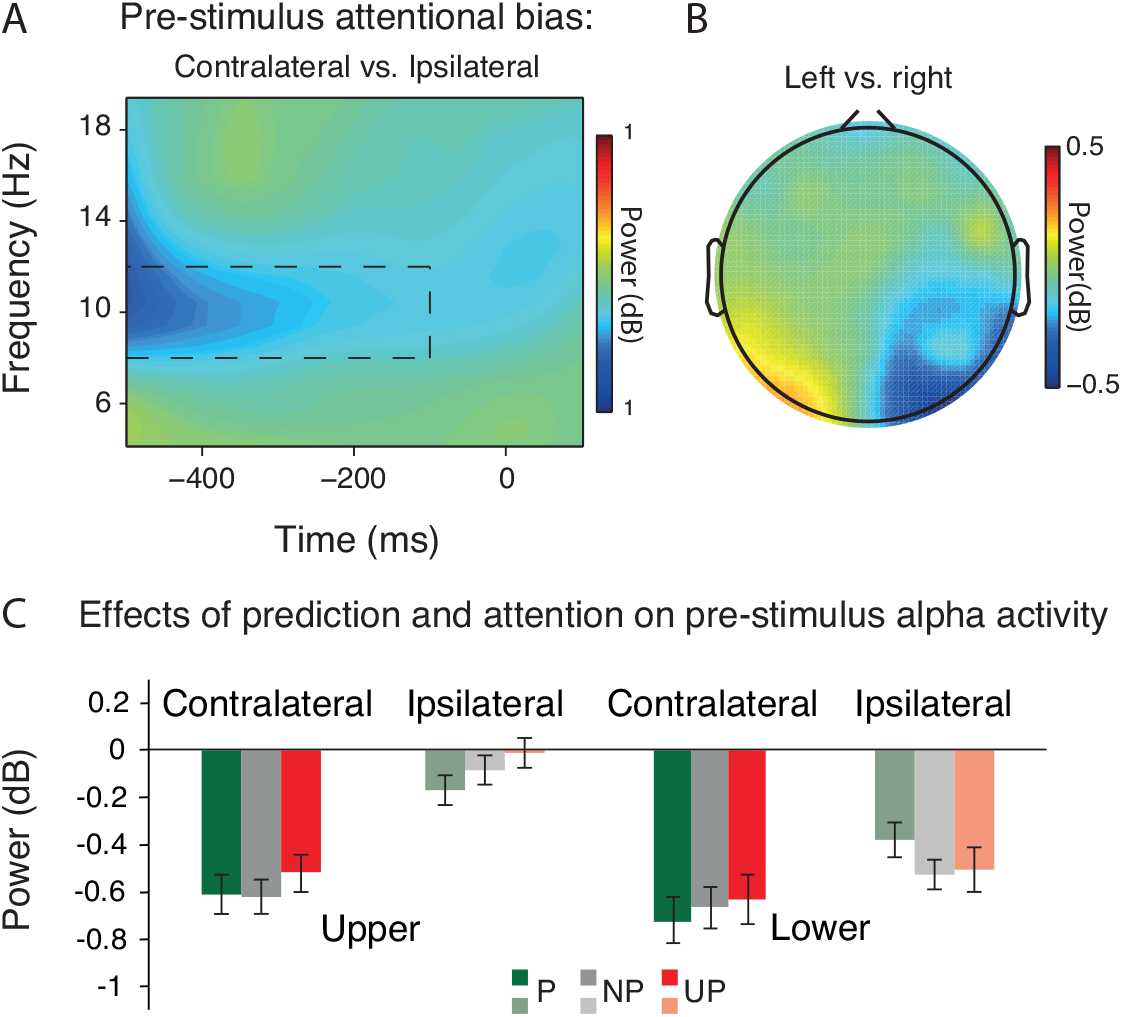
Effects of top-down attention and prediction on pre-stimulus alpha power. (**A**) Shown are differences in average power values across time as a function of frequency at electrodes contralateral vs. ipsilateral to the cued (attended) stimulus location (PO3/7 and PO4/8). Attention (collapsed over prediction conditions and upper and lower locations) was associated with reduced pre-stimulus (m500 to −100ms) alpha power (8-12 Hz) (marked by the black dashed rectangle) over contra-compared to ipsilateral posterior scalp regions. (**B**) The scalp distribution of pre-stimulus alpha power in a −500 to −100ms window over posterior sites as a function of cue direction: left vs. right (collapsed over upper and lower locations). As can be seen, spatial attention was associated with a lateralization of pre-stimulus alpha activity over lateral posterior scalp regions. (**C**) Prediction modulated the effect of attention on pre-stimulus alpha lateralization (i.e., the magnitude of pre-stimulus alpha power at electrodes contralateral vs. ipsilateral to the attended stimulus location (PO3/7 and PO4/8)), but only when locations in the lower field were attended. Specifically, alpha power was only higher over ipsilateral than over sites contralateral to predicted lower-field locations.

#### P1 and N1 modulations

Given the lack of a C1 modulation, it is important to demonstrate that later effects of attention and prediction on visual processing shown consistently in previous research (Di Russo et al., 2003; Marzecová et al., 2017), were replicated. In line with this body of research (e.g., Hillyard, Vogel, & Luck, 1998) and as can be seen in Figure 6, the P1 was enhanced by attention, although specifically over contralateral posterior scalp regions (contralateral: t_19_=3.103, p=.006; ipsilateral: t_19_=-0.182, p=.858), as reflected by a significant interaction between Attention and Hemisphere (F_1,19_=7.221, p=.015; main effect of Attention: F_1,19_= 1.395, p=.252). The P1 was also modulated by predictions over contralateral posterior regions, albeit only marginally (Hemisphere x Prediction interaction: F_2,18_=3.184, p=.065; main effect Prediction: F_2,18_= 1.017, p=.330). The effects of attention and prediction on the P1 were independent of each other, as reflected by a non significant interaction between Attention and Prediction (F_2,18_=.317, p=.732). All other interaction effects of Attention and/or Prediction with Field and/or Hemisphere were nonsignificant (all p’s >.103).

As expected, attention also enhanced the amplitude of the N1 component (F_1,19_=22.431, p<.001; Fig. 6). This attention effect was bilateral (Attention x Hemisphere: F_2,18_=0.272, p=.608) and observed independent of stimulus location (Attention x Field: F_2,18_=0.006, p=.941), although it was somewhat larger for lower field stimuli over the ipsilateral hemisphere, as indicated by a significant Attention x Hemisphere x Field interaction (F_1,19_=7.843, p=.011) (see Fig. 6).

**Figure 6.**
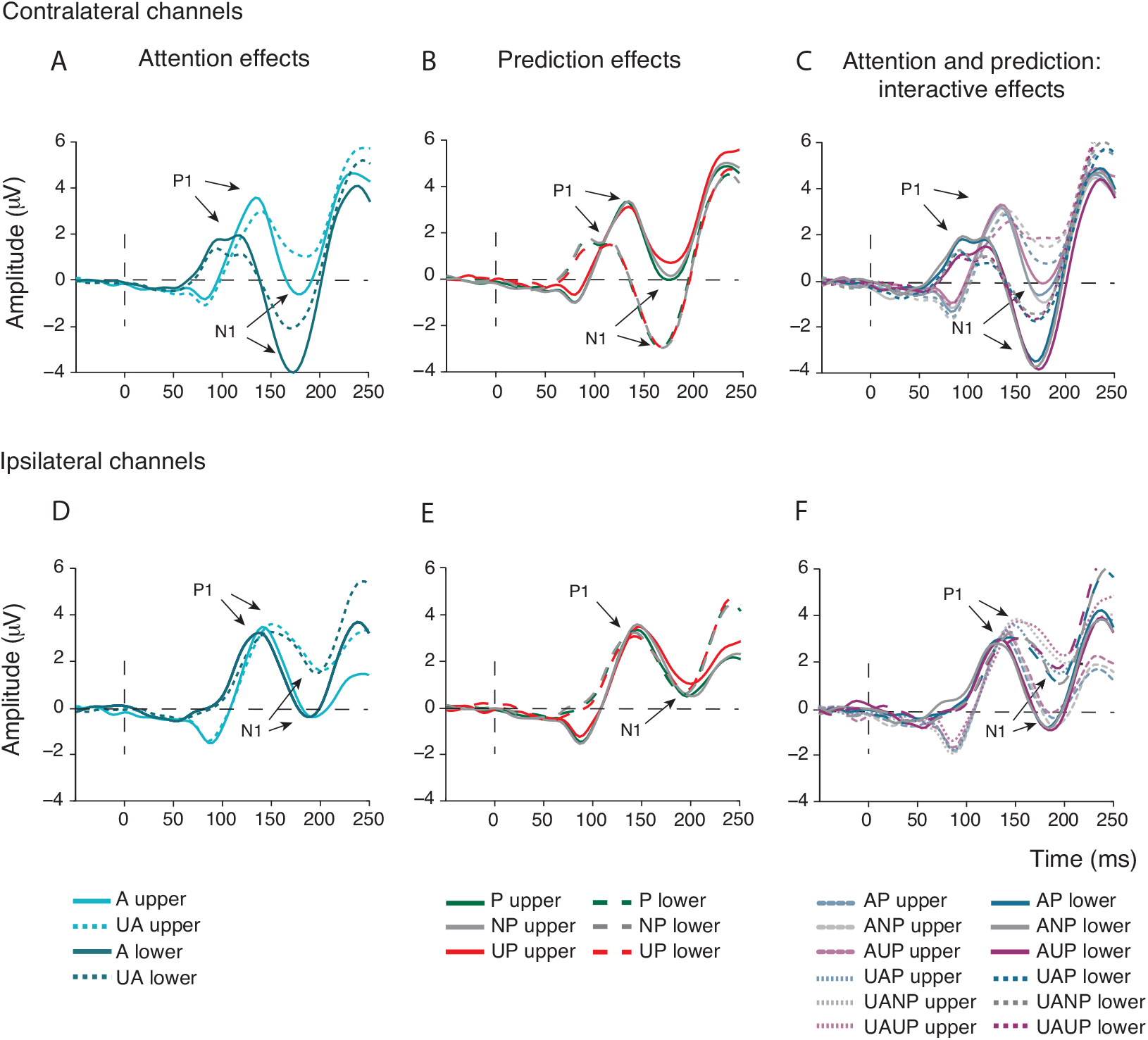
Effects of attention and prediction on later stages of visual information processing indexed by the P1 and N1 ERP components. (**A, D**) Attention was also associated with a larger contralateral P1 and bilateral N1. (**B, E**) Predictions did not modulate P1 and N1 amplitudes. (**C, F**) Prediction did marginally modulate the N1 peak in interaction with attention.

Prediction did not modulate the N1 (F_2,18_=1.493, p=.251), but a marginally significant interaction between Prediction x Field (F_2,18_=2.141, p=.063) suggested that the differences in N1 amplitude between prediction conditions might be greater at upper locations (see Fig. 6). Notably, the interaction between Attention and Prediction (F_2,18_=3.311, p=.06) was marginally significant. With the exception of a 4-way interaction between Attention, Prediction, Field and Hemifield (F_2,18_=2.966, p=.038), suggesting that the interaction between attention and prediction differed significantly across specific combinations of stimulus locations and hemisphere, none of the other interaction effects of Attention and/or Prediction with Field and/or Hemisphere reached significance (all p’s>.077).

Thus, attention robustly modulated later visual processing, as indexed by the P1 and N1, whereas effects of prediction on later visual processing were relatively weak and only marginally significant.

#### P3a and P3b modulations

Prediction and attention have also robustly been shown to modulate later, post-perceptual stages of information processing (Friedman et al., 2001; Polich, 2007). In line with these observations, attention and prediction, independently and in interaction, modulated the amplitude of the P3a and P3b ERP components. Both attention (F_1,19_=29.040, p<.001) and prediction (F_2,18_= 18.330, p<.001) reduced the amplitude of the P3a component (Fig. 7). Moreover, as suggested by a significant interaction between Attention and Prediction (F_2,18_=8.831, p<.001), the effect of prediction was larger in the unattended compared to the attended condition (Fig. 7C).

**Figure 7.**
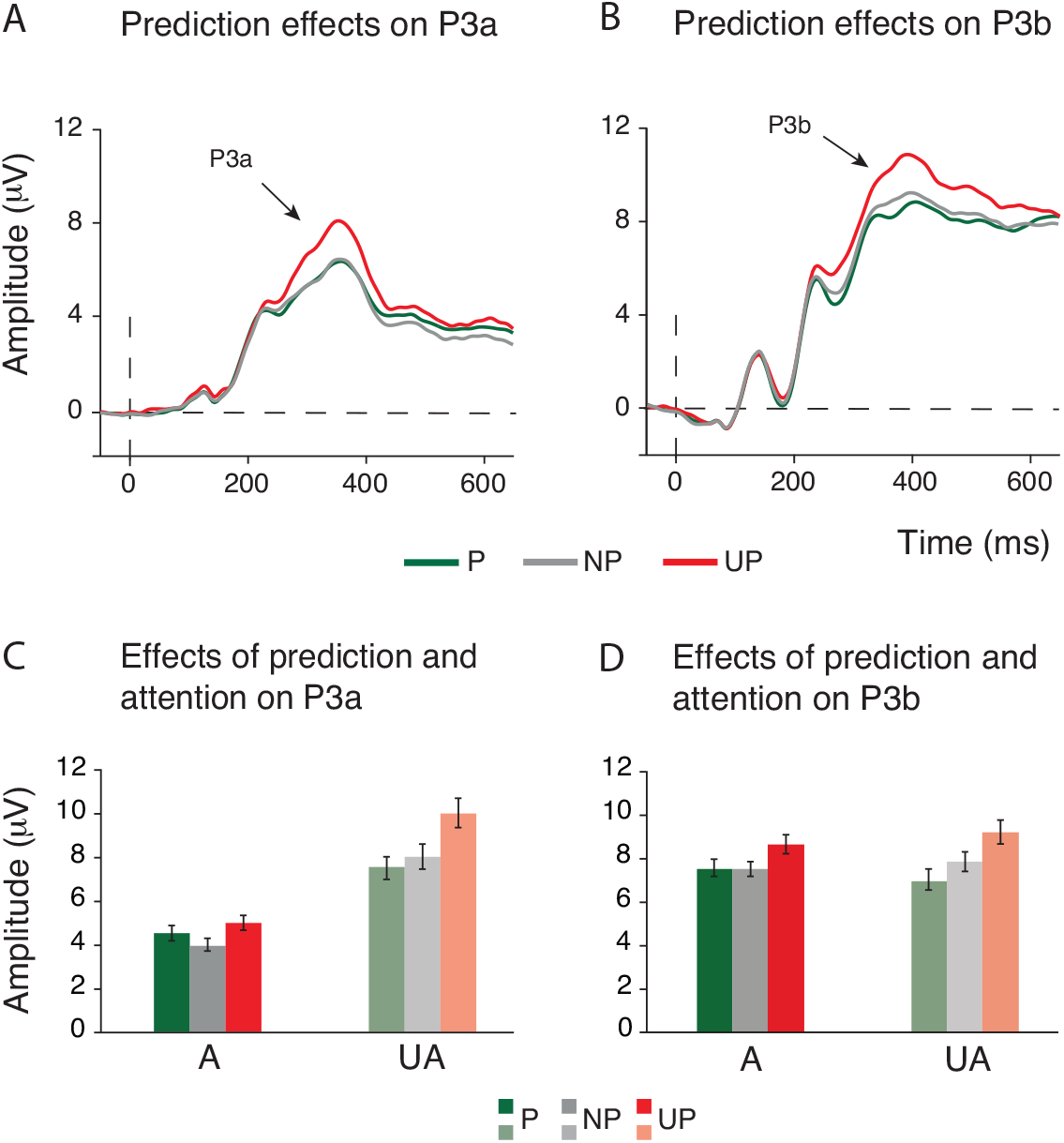
Effects of attention and prediction on post-perceptual stages of information processing as indexed by the P3a and P3b components. (**A**) Prediction modulated the size of the P3a novelty response (~300-400ms post-stimulus) in the expected way: the P3a was higher to unpredicted (UP) compared to predicted (P) and non-predicted (NP) stimuli. (**B**) Prediction also modulated the size of the P3b component (~340-440ms post-stimulus), such that the amplitude of the P3b was inversely related to the stimulus probability. (**C**) Post-perceptual effects of prediction also depended on attention. The P3a response was highest to unpredicted stimuli at both attended and unattended locations. (**D**) The amplitude of the P3b response scaled inversely with stimulus predictability at attended and unattended locations. The largest P3b response was again observed to unpredicted stimuli, however, non-predicted stimuli elicited a smaller P3b than predicted stimuli.

The P3b was also modulated by predictions, but in contrast to the P3a, not by attention, as reflected by a significant main effect of Prediction (F_2,18_=32.283, p<.001) and an insignificant main effect of Attention (F_1,19_=.058, p=.812). As expected, and can be seen in Figure 7B, the amplitude of the P3b component scaled down as the probability of a stimulus increased. The interaction between attention and prediction was not significant (F_2,18_=4.474, p=.199), suggesting that prediction effects on the P3b were not modulated by attention.

To summarize the results of Experiment 1, while ERP and multivariate analyses did not provide any evidence for a modulation of visual activity before 80ms post-stimulus by prediction or attention, these top-down factors were associated with robust modulations of later stages of information processing. Moreover, modulations of pre-stimulus activity indicated that prediction and attention biased sensory processing in advance. Nevertheless, this did not affect the very first stage of cortical information processing.

#### Eye-tracking

The analysis of eye tracking data indicated that the mean eye position at the time of stimulus presentation (−200 to 100ms) was not modulated by Prediction in the x (F_2,18_=.443 p<.714) or y (F_2,18_=1.405 p<.271) direction. Yet, attention did affect mean eye position at the time of stimulus presentation, as indicated by a significant main effect of Attended Location in both directions (x direction: F_1,19_=83.254 p<.001; y direction: F_1,19_=56.861 p<.001). Yet, post-hoc inspection of the data suggested that, albeit consistent across subjects, these differences in eye position were very minor in magnitude (the average difference in gaze direction between cued locations in x and y directions was 0.16° and 0.4°, respectively; absolute average deviations from fixation were: x direction attend low (right): 0.06°; x direction attend up (left): −0.1°; y direction attend low (right): 0.24°; y direction attend up (left): 0.16°). It is unlikely that such minor differences can explain possible differences in stimulus-evoked ERPs between attended and unattended stimuli, which is supported by the lack of an attentional modulation of the first ERP component, the C1 (reported above). Moreover, the interaction between Attended Location and Prediction was not significant (x direction: F_1,19_=2.018, p=.162; y direction: F_1,19_=.396 p=.679).

### Experiment 2

In Experiment 1 we obtained moderate to strong evidence against a modulation of stimulus-driven activity by spatial prediction and attention before 80ms. While our C1 peak analysis suggested that prediction may modulate the later phase of the C1, this effect was statistically weak, not supported by Bayesian analyses, and in a direction opposite to what one would expect based on predictive processing theories in which predictions are proposed to reduce visual responses (i.e., prediction errors) (e.g. Alink et al., 2010; Friston, 2009). To more conclusively establish the absence of top-down effects before 80ms, we conducted a second EEG experiment to determine if our findings would replicate using a design that was further optimized to detect activity generated by V1. We again independently modulated spatial attention and prediction using a similar cueing paradigm as in Experiment 1 with three important improvements. First, we used large-scale, high-contrast texture stimuli that are known to elicit large C1’s (e.g., see Pourtois, Rauss, Vuilleumier, & Schwartz, 2008). It is possible that in Experiment 1, we did not observe any attention- and/or prediction-based modulations of the earliest stage of visual information processing due to the relatively low signal-to-noise ratio at this stage of processing. By boosting the strength of the bottom-up input to V1, we expected that our sensitivity to measuring weak top-down effects would be improved (Slagter et al., 2018). As a second improvement, in Experiment 2, we used a task in which participants had to perform an orientation discrimination task (vs. a ring detection task), which conceivably relies on V1 activity. Third and lastly, we presented stimuli in the left or right upper visual field (versus at two diagonally opposite locations, one in the upper and one in the lower visual field, as in Experiment 1). By presenting stimuli only in the upper visual field, we importantly aimed to minimize overlap between the C1 and the subsequent P1 component (Qu & Ding, 2018), which is only problematic for lower-field stimulation. This allowed us to better isolate possible C1 effects. In addition, some studies report that modulations of the C1, e.g., by attentional load (Rauss, Pourtois, Vuilleumier, & Schwartz, 2009) and perceptual learning (Pourtois, Rauss, Vuilleumier, & Schwartz, 2008), are only present for upper visual field stimuli (see also Slotnick, 2018). Thus, in Experiment 2 we aimed to determine if we could observe early (before 80ms) effects of prediction and/or attention when using V1-tuned stimuli, an orientation discrimination task, and an upper-field stimulation protocol, or would replicate our null results from Experiment 1.

#### Behavior

Contrary to the observed prediction-related increase in reaction time in Experiment 1 and in Kok et al. (2012), stimulus predictability had no influence on average reaction time to target stimuli (F_2,11_=0.819; p=.632, P=528.7, SD=108.2; NP=531.6, SD=109; UP=535, SD=100.4) (see Fig. 8). Yet, as in Experiment 1 and Kok et al. (2012), we found that stimulus predictability did not affect response accuracy (F_2,11_=1.550, p=.225, P=74.6, SD=8.2; NP=76.2, SD=8.1; UP=76.6, SD=7.3) or sensitivity to target signals, expressed in d’ (F_2,11_=.150, p=.863; d’ P=1.58, SD=0.3; d’ NP=1.6, SD=0.3; d’ UP=1.61, SD=0.44). While the mean accuracy was comparable between the experiments, reaction times were about 60-100ms slower in Experiment 2. Moreover, d’s were consistently lower in Experiment 2. These results indicate that participants took more time to respond and were less sensitive to target signals in Experiment 2, indicating that our task design changes made the task used in Experiment 2 more difficult. It is unclear which changes specifically may have led to these differential results between experiments (e.g., the use of a non-spatial symbolic cue in Experiment 2 vs. a spatial cue in Experiment 1, the line element stimuli in Experiment 2 vs. the Gabor stimuli in Experiment 2).

**Figure 8.**
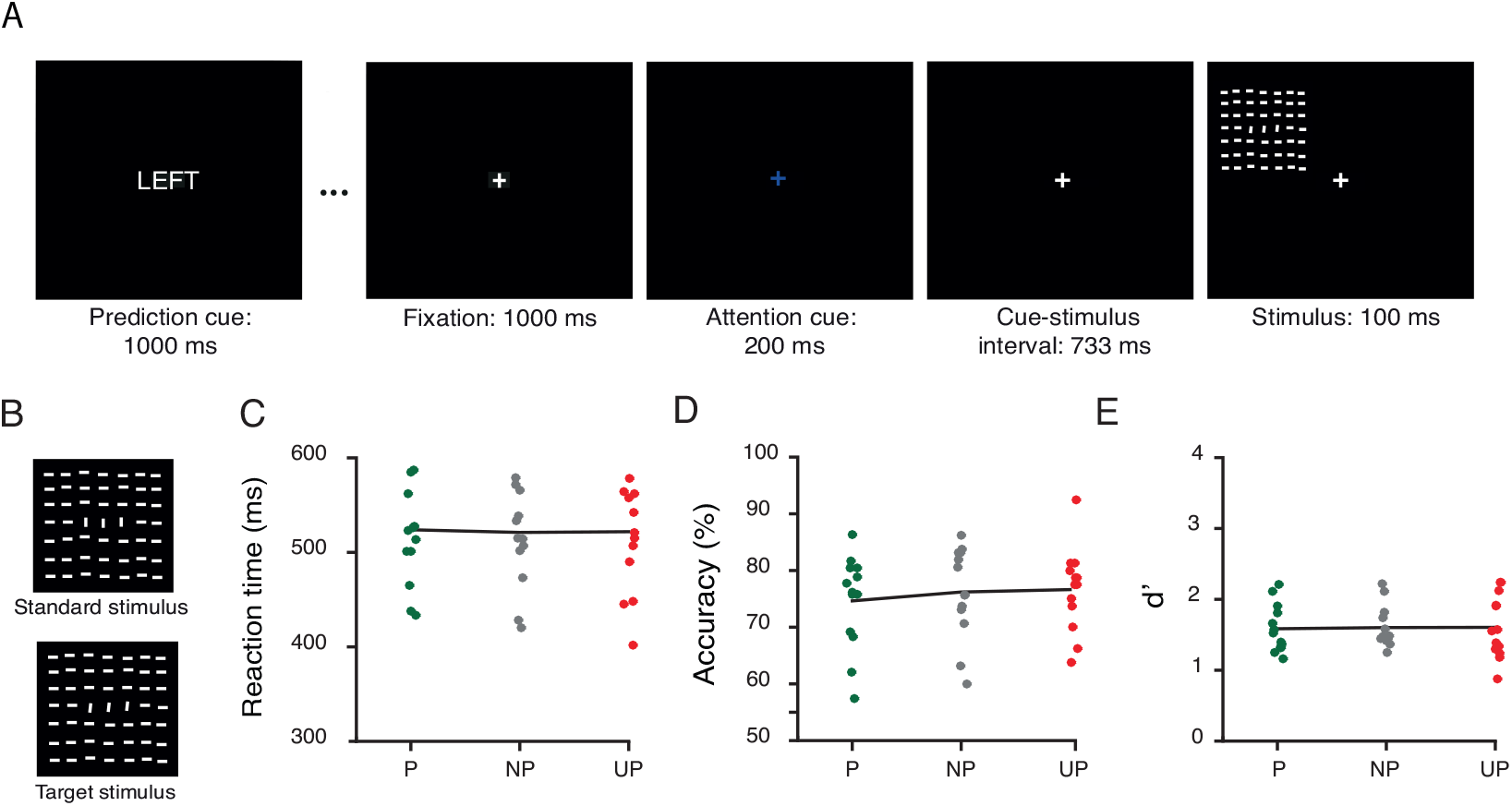
Stimuli, experimental task and behavioral results of Experiment 2. (**A**) Example of a trial of the spatial cueing task. Each block of 20 trials started with the presentation of a prediction cue (the word “LEFT”, “RIGHT’” or “NEUTRAL”) signaling the likelihood that a stimulus would occur at the upper left or right location in that block. Each trial started with the presentation of an attention-directing cue, which instructed participants to covertly direct their attention to the cued location (centrally presented fixation cross in red or blue signaling right or left location, respectively). After a fixed interval, the cue was followed by a stimulus, a texture stimulus, at the cued or uncued location. Participants had to press the left mouse button in case of a target stimulus at the cued location. The depicted sequence shows an example of a predicted attended trial in which a target stimulus is presented at the more likely and relevant location. (**B**) Target stimuli appeared on 25% of trials and were texture stimuli with the foreground region (three vertically oriented lines in the center of a stimulus) tilted towards the left or right with respect to the foreground region. On a standard stimulus, the foreground region formed a 90° degrees angle with respect to the background lines. There was no behavioral benefit of stimulus predictability on reaction times (**C**), accuracy (**D**) or d prime (**E**). P: predicted; NP: non-predicted; UP: unpredicted.

#### ERP results

In Experiment 2, we examined attention/prediction effects on the early (50-80ms) and late phase of the C1 and on early patterns of activity, to address our main question of whether top-down factors can modulate initial afferent activity and representational content before ~80ms.

**Figure 9.**
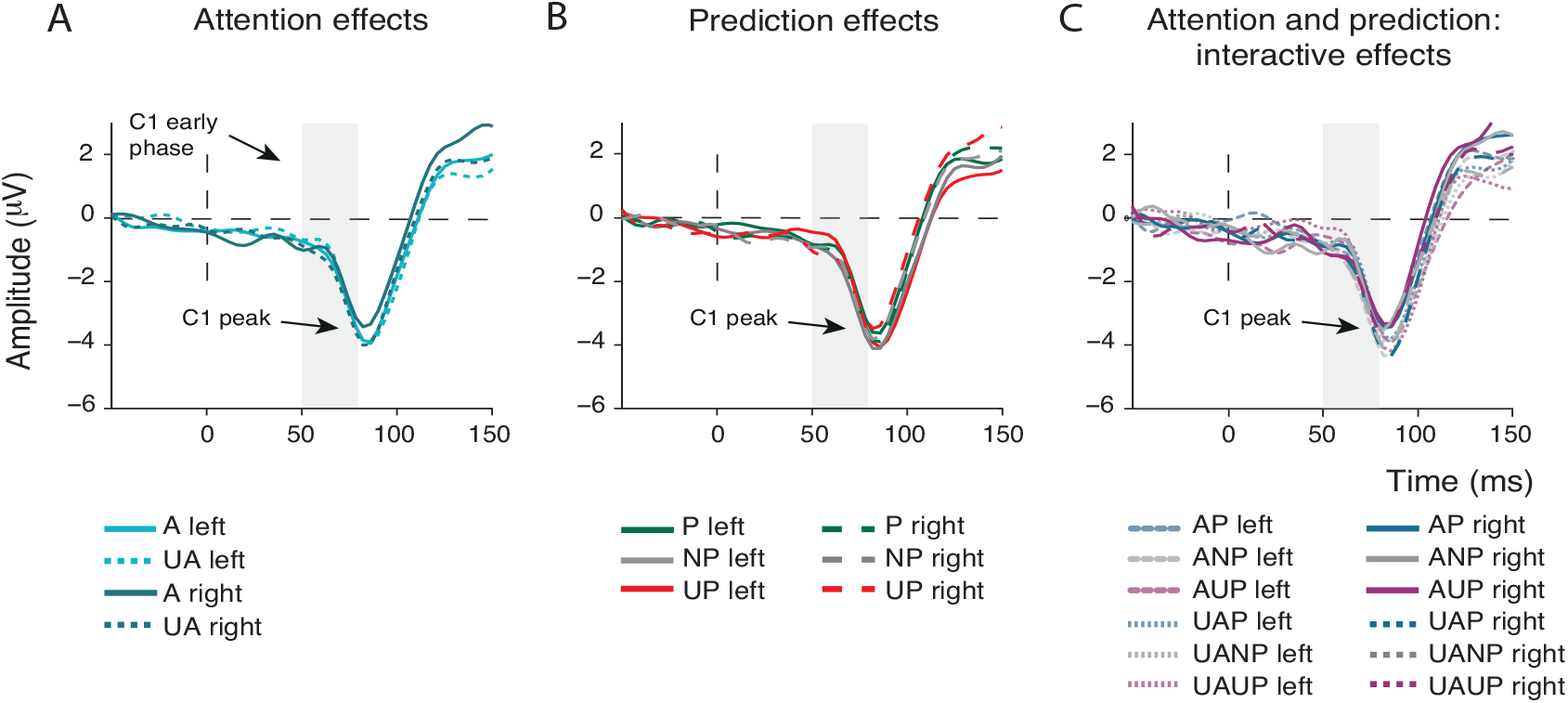
Effects of attention and prediction on the first feedforward sweep of cortical information processing in Experiment 2. (**A**) Attention did not modulate C1 amplitude (neither in the early time window, nor at the peak). (**B**) Prediction also did not modulate the C1 (neither in the early time window nor at the peak). (**C**) Prediction and attention in interaction did not modulate the amplitude of the C1 (neither in the early time window nor at the peak). A: attended; UA: unattended; P: predicted; NP: non-predicted; UP: unpredicted.

As expected, the texture stimuli used in Experiment 2 elicited a C1 that was on average almost twice as large as in Experiment 1. Nevertheless, replicating the C1 results from Experiment 1, analyses examining effects of prediction and attention did not reveal any evidence for top-down modulation of C1 amplitude during the early phase (50-80ms poststimulus) or the late phase (peak) of the C1 in Experiment 2. Specifically, during the early phase of the C1, the main effects of Attention (F_1,12_ = .078, p=.785) and Prediction (F_2,11_=2.495, p=.318), as well as their interaction (F_2.11_ =0.001, p=.999), were all far from significance. This was supported by Bayesian analyses, which showed substantial evidence for the null hypothesis against the main effects of Attention (BF_01_=4.95), Prediction (BF_01_=6.99) and their interaction (BFinclusion =8.33). The same was true for the late phase of the C1 (main effect of Attention: F_1,12_= 1.083, p=.318; main effect of Prediction: F_2,11_=2.405, p=.376; interaction Attention and Prediction: F_2,11_=.522, p=.607). This was further corroborated by Bayesian statistics, which again provided substantial support for the absence of these effects (Attention BF01=3.09; Prediction BF01=8.55; Attention x Prediction BF_inclusion_=7.7). Thus, in Experiment 2, we again found no evidence for a modulation of the first feedforward sweep of cortical activity by either prediction or attention.

#### MVPA results

In line with the ERP findings and the multivariate results from Experiment 1, decoding analyses did not reveal modulations of the pattern of EEG activity by attention or prediction before 80ms (attended vs. unattended: 164 and 852ms, p=.002; prediction: 414 and 508ms, p=.015). These overall smaller effects compared to experiment 2 probably have to do with the smaller sample size affecting the statistical power of the performed tests.

#### Eye-tracking

For only 5 of our 13 participants we could collect eye-tracking data in Experiment 2. For those participants, we compared average deviations in x and y direction. The only significant effect observed was a main effect of Attended Location for deviations in x direction (x direction: F_1,12_=6.551, p=.025; y direction: F_1,12_=0.512 p=.488). Similarly to what we found in Experiment 1, the difference in the eye position in the x direction after attention cues pointing to the left versus to the right was very small: 0.24°. Moreover, average deviations from fixation (x direction attend right: 0.05°; × direction attend left: −0.19°; y direction attend right: 0.06°; y direction attend left: 0.05°) were very small. Thus, these eye tracking data also suggest negligible differences in eye position as a function of attention. Importantly, as in Experiment 1, the main effect of Predicted Location and the interaction effect between Predicted Location and Attended Location were non-significant (all p’s>.168).

For participants without the eye tracking data (n=8), we relied on averaged HEOG voltage values to determine that fixation was maintained at the time of stimulus presentation. A repeated measures ANOVA showed that the main effects of Attended Location, Predicted Location and their interaction were not significant (Attended Location: F_1,7_=0.00, p=.994; Predicted Location: F_2,6_=0.141, p=.898; Attended Location x Predicted Location: F_2,6_ =0.404, p=.684), suggesting that there was no systematic bias in the average gaze in favor of relevant and/or predicted locations. In line with this, the average deviation of the HEOG amplitude after a cue pointing to the left was −0.05 μV, and −0.27 μV after a cue pointing to the right location in the upper visual field, which corresponds approximately to average eye deviations of 0.005° to the left and 0.024° to the right (see Mangun & Hillyard, 1991). These results suggest that differences in eye position at the time of stimulus presentation between conditions likely did not affect our ERP and MVPA results.

## Discussion

Influential predictive processing theories postulate that predictions derived from past experience can influence sensory information processing across the cortical hierarchy, and that attention can modulate these effects by boosting the precision of predictions (Friston, 2009; Kok, Rahnev, et al., 2012; Rao, 2005). In two studies, we exploited the high temporal resolution of EEG, with an optimized design to detect activity generated by V1. However, we found no evidence that spatial predictions (stimulus likelihood), either independently or in interaction with attention (stimulus relevance), modulated the earliest stage of cortical visual information processing indexed by the early phase of the C1 (50-80ms post-stimulus). Strikingly, we did observe modulations of pre-stimulus alpha-band oscillatory activity, suggestive of the implementation of a top-down sensory bias prior to stimulus presentation. Nevertheless, this was not accompanied by a modulation of the earliest visual-evoked activity. These findings extend previous results from attention studies by showing that visual activity prior to ~100ms may be impenetrable to top-down influences in general (Baumgartner et al., 2018; Di Russo et al., 2012, 2003; Martinez et al., 2001; Martínez et al., 1999; Noesselt et al., 2002). Replicating previous findings (e.g., Baumgartner et al., 2018; Marzecová et al., 2017), attention and prediction did affect subsequent stages of information processing, as reflected in modulations of the P1, N1 and P3 ERP components. These ERP results were corroborated by MVPA analyses, which revealed the earliest attentional modulations from ~ 130ms and prediction modulations from ~240ms after stimulus presentation.

In previous studies that found no effect of attention on the C1, attention-directing cues had no predictive value (Baumgartner et al., 2018; Di Russo et al., 2003; Simon P. Kelly et al., 2008), which may have affected the level of attention prior to stimulus presentation and thereby how early attention influenced visual stimulus processing. However, our results suggest that even when attended stimuli are highly likely, the earliest stage of visual information processing may remain unaffected by top-down influences. One study did report C1 modulations by attention, despite the equal stimulus likelihood at attended and unattended locations (Kelly et al. (2008). Yet, following a highly similar experimental and analytical protocol, we and others (Baumgartner et al. 2018), in a larger sample size, did not replicate this original finding. At present, one can only speculate as to what may cause these differential effects (Kelly & Mohr, 2018; Slotnick, 2018). It is possible that small differences in the stimuli used in our Experiment 1 and in Baumgarter et al. (2008) versus Kelly et al. (2008) contributed to the discrepancy in findings (see Kelly & Mohr, 2018). However, using a V1-tuned task and stimuli, in Experiment 2, we still obtained no evidence for a top-down modulation of the C1. Effects of spatial attention on the C1 were also reported in a study by Rauss et al. (2009). In that study, the authors varied attentional load at fixation and found a decrease in C1 amplitude to stimuli presented in the periphery when load was high. Yet, as effects of spatial attention in the periphery were inferred only indirectly - in relation to the central load manipulation, and eye tracking was not used, alternative explanations for the diminished C1 cannot be excluded (e.g., differences in pupil size or eye-movements between load conditions, Bombeke, Duthoo, Mueller, Hopf, & Boehler, 2016). In general, the five EEG/MEG studies in humans (Kelly et al., 2008; Poghosyan & Ioannides, 2008; Rauss et al., 2012; Rauss et al., 2009; Slotnick et al., 2002) that reported modulations of the C1 component by spatial attention were either not replicated (Rauss et al., 2009; see Rauss et al., 2012 for opposite results, and Ding, Martínez, Qu, & Hillyard, 2014 for a null replication), or suffer from methodological issues, such as low trial numbers (Poghosyan & Ioannides, 2008) or uncontrolled eye-movements (Slotnick et al., 2002; see Baumgartner et al., 2018 for a more extensive review of some of these studies), which calls for caution and replication.

Albeit replicated across two experiments, the present null finding cannot solely be taken to discard the notion from hierarchical predictive processing models of vision that predictions are implemented as early as V1 (Clark, 2013; Friston, 2009). It is possible that despite our optimal design to detect V1 activity, prediction effects in V1 were too weak to be detected at the scalp level, or take longer to be established than was allowed for in our design. Yet, we orthogonally manipulated prediction and attention in a similar manner as a previous fMRI study, which observed joint modulation of neural responses in V1 by these top-down factors (Kok, Rahnev, et al., 2012). Moreover, in Experiment 1, we replicated the behavioral effect of predictions on response speed reported in this fMRI study, indicating that stimulus location likelihood was adaptively tracked.

Another striking aspect of the present findings is that even though the initial phase of the C1 was not modulated by prediction/attention, we did observe attention-related effects on pre-stimulus posterior alpha-band activity, indicative of the establishment of a top-down visual bias (Horschig et al., 2014; Kelly et al., 2006; Sauseng et al., 2005; Worden et al., 2000). One possible explanation to reconcile these opposing findings could be that prediction signals and feedforward stimulus-driven activation recruit distinct neuronal populations (Bastos et al., 2012; Kok, Bains, Mourik, Norris, & Lange, 2016). Another, not mutually exclusive, explanation could be that predictions selectively modulate stimulus-driven activity at higher frequencies, specifically in the gamma range, that are not captured in ERPs and are in general difficult to measure with scalp EEG. Higher cortical regions implement top-down predictions in hierarchically lower cortical regions though synchronization of activity in lower-frequency ranges (i.e., the alpha/beta band), whereas prediction violations are propagated from lower to higher cortical regions through synchronization of gamma band activity (Arnal & Giraud, 2012; Michalareas et al., 2016). Indeed, a recent MEG study found that invalid predictions increased gamma activity induced by task-relevant stimuli in V1, however, not until 130ms after stimulus presentation (Auksztulewicz, Friston, & Nobre, 2017). Another MEG study employing a probabilistic cuing task also reported stimulus-induced increases in gamma-band activity by attention, which, in contrast to Auksztulewisz et al., decreased as a function of stimulus predictability (Bauer, Stenner, Friston, & Dolan, 2014). Yet, there too, these gamma modulations occurred after 100ms. Thus, top-down modulations of high-frequency gamma activity, like our ERP effects, also seem to occur after the first feedforward sweep of information processing. Nevertheless, an important avenue for future studies is to further determine the role of neural oscillations in predictive processing.

Overall, our findings corroborate the so-called “majority view” (Slotnick, 2013), according to which attention can only bias processing in V1 through delayed feedback from extrastriate visual areas. This view is based on human EEG studies of attention (e.g., Di Russo et al., 2003, 2012; Martinez et al., 2001; Martínez et al., 1999; Noesselt et al., 2002), but also supported by monkey studies, in which recordings were made directly from V1 neurons. Also in these studies, typically, no attention effects are observed on neural activity in V1 prior to 100ms post-stimulus, and/or no V1 effects are observed at all (Briggs, Mangun, & Usrey, 2013; Roelfsema, Tolboom, & Khayat, 2007; Sharma et al., 2014; Stănişor, Togt, Pennartz, & Roelfsema, 2013). All together, these observations suggest that predictions regarding visual input may not (necessarily) influence the first cortical processing stage. This also suggest that previous observations based on the sluggish fMRI BOLD response, showing that predictions can modulate BOLD responses in V1 (Alink et al., 2010; Kok, Jehee, et al., 2012; Kok, Rahnev, et al., 2012; St. John-Saaltink, Utzerath, Kok, Lau, & de Lange, 2015), conceivably do not reflect a modulation of the first feed-forward sweep of cortical information processing, but later, recurrent effects. Indeed, in our study, both prediction and attention modulated later stages of information processing. Prediction effects on the P1 were admittedly weak and absent for the N1 (see Lasaponara et al., 2017 for similar findings), but prediction effects on the P3a and P3b were robust and exhibited a pattern that is consistent with earlier work demonstrating their role in novelty processing and prediction updating (Friedman et al., 2001; Marzecová et al., 2017; Polich, 2007). Consistent with predictive processing accounts and the idea of the inverse scaling of neural response in relation to the size of prediction errors (den Ouden, Kok, & de Lange, 2012; Friston, 2009; Hohwy, 2012), we observed larger P3a and P3b responses to unpredicted than to predicted stimuli at both attended and unattended locations.

In conclusion, we found no evidence that prediction and attention, independently or in interaction, modulated neural activity prior to 80ms after stimulus presentation, even though pre-stimulus activity indicated the establishment of a top-down visual bias. This conclusion converges with evidence from a large body of research on attention. The results presented here additionally indicate that the absence of top-down modulation of visual afferent activity by spatial attention in previous studies cannot be explained by dampening of attentional effects due to equal stimulus likelihood at attended and unattended locations. Overall, prediction and attention may modulate V1 processing through delayed feedback from extrastriate visual areas, or, alternatively, through mechanisms that are not captured by M/EEG.

## Acknowledgements

We thank Simon Kelly for sharing his task code and stimuli with us for Experiment 1 and feedback on an earlier version of this paper. We also thank Floris de Lange for useful input on the task design of Experiment 1. We are grateful to Johannes Fahrenfort for his assistance with the multivariate EEG analyses, and to Gilles Pourtois for his advice regarding the visual stimuli used in Experiment 2. We also thank Tara van Viegen for her help in pilot work that led to this study. This work was supported by an ERC grant to H.A.S. by the H2020 European Research Council [ERC-2015-STG-679399].

